# Sustained alpha oscillations serve attentional prioritization in working memory, not maintenance

**DOI:** 10.1101/2025.10.16.682668

**Authors:** Yuanyuan Weng, Jelmer P. Borst, Elkan G. Akyürek

## Abstract

Recent theory on the neural basis of working memory (WM) has attributed an important role to “activity-silent” or -quiescent mechanisms, suggesting that sustained neural activity might not be essential in the retention of information. This idea has been challenged by reports of ongoing neural activity in the alpha band during WM maintenance, however. The precise role of these alpha oscillations is unclear: Do they reflect attentional prioritization of stored information, or do they serve as a general maintenance mechanism, for instance to periodically refresh synaptic traces? To address this, we designed a visual WM task involving two memory items, one of which was prioritized by being tested first for recall. The task included both short (1 s) and a long (3 s) delay intervals between encoding and retrieval. The long delay condition allowed us to test whether the alpha-based decoding effects persist beyond the early delay period, thereby putting accounts that attribute alpha activity to generic maintenance processes to the test. Time-resolved decoding analyses revealed that both tested-first and tested-second items were initially decodable following stimulus presentation. However, only the tested-first item exhibited sustained decodability throughout the delay, particularly in the long delay condition, where it transitioned into a stable coding scheme. This prolonged representation was selectively supported by induced alpha power, which reliably tracked the prioritized tested-first item, but not the deprioritized tested-second one. Impulse-based decoding further confirmed this asymmetry, showing a selective increase in readout for the tested-second item only when it became immediately task-relevant. Together, these findings suggest that sustained alpha-band activity primarily reflects attentional prioritization, rather than general memory maintenance. Unattended, deprioritized items appear to transition into an activity-quiescent state, consistent with models of synaptic storage in WM.

## 1. Introduction

Working memory (WM) is a crucial cognitive system that maintains and stores information, guiding thought and behavior in the absence of direct sensory input (Baddeley, 2003). There has been growing interest in understanding its underlying neural mechanisms, particularly regarding how neural activity sustains information during the retention periods. The traditional view is that WM retention depends on persistent activity, wherein sustained neural firing preserves representations of memory items across the delay periods (Constantinidis et al., 2018; Goldman-Rakic, 1995; Kojima & Goldman-Rakic, 1982), ensuring continuous availability of stored information. However, persistent activity has not consistently been observed during WM tasks. An alternative account is based on “activity-silent” ^1^ or - quiescent neural mechanisms (Stokes, 2015; Wolff et al., 2017), which proposes that information can (also) be maintained through short-term synaptic changes (Barak & Tsodyks, 2014; Buonomano & Maass, 2009; Choudhary et al., 2024; Kozachkov et al., 2022; Lundqvist et al., 2018; Miller et al., 2018; Mongillo et al., 2008; Pals et al., 2020; Panichello et al., 2024; Spaak & Wolff, 2025; Sprague et al., 2016). In this view, synaptic changes serve as a latent memory trace that can still be rapidly accessed and made behaviorally available when the stored information becomes relevant. Recent computational evidence supports the idea of activity-quiescent WM, demonstrating that short-term synaptic plasticity (STSP), modelled by calcium-dependent synaptic facilitation, effectively maintains memory representations without sustained neural firing (Pals et al., 2020).

Empirical research also seemed to provide evidence for the activity-quiescent framework, at least initially. For example, Wolff et al. (2017) presented two memory items, instructing participants to prioritize only one of them for the upcoming test, thereby deprioritizing the unattended item. Using a functional perturbation approach involving a high-contrast, task-irrelevant visual impulse during the maintenance period, they found that WM contents remained decodable from the neural impulse response, even in the absence of sustained delay-period activity (see also Rose et al., 2016). A key finding of this study was that unattended memory items that were deprioritized after a retrospective cue exhibited little to no sustained neural activity, suggesting they might be stored in an activity-quiescent state rather than through continuous neural firing. However, a later re-analysis, in which the same EEG data were examined using alpha-band (8-12 Hz) power rather than raw EEG voltage, challenged this interpretation (Barbosa et al., 2021). In contrast to previous reports (LaRocque et al., 2013; Rose et al., 2016), Barbosa and colleagues (2021) showed that unattended items were decodable from alpha power during the delay period, although not consistently so throughout the entire delay interval. These outcomes suggest that maintenance in WM might be supported by oscillatory activity, which has been proposed by several authors (F.-W. Chen et al., 2023; Foster et al., 2017; Fulvio et al., 2024; Sutterer et al., 2019; ten Oever et al., 2020).

Alpha-band oscillations in particular may be crucial in WM, as they have been implicated in regulating attentional processes, inhibiting irrelevant information, and potentially influencing memory retention (Bonnefond & Jensen, 2012; X. Chen et al., 2023; Klimesch, 2012; Vries et al., 2020). Research also indicates that alpha power increases during WM retention, particularly in parietal and occipital regions, which may reflect active involvement in WM storage (Jensen et al., 2002). However, the exact function of alpha oscillations in WM maintenance remains unclear. Do alpha-band oscillations primarily reflect attentional prioritization mechanisms acting on WM representations (Bae & Luck, 2018), or are they more closely related to more general maintenance mechanism-related processes, sustaining memoranda, independent of prioritization (cf. Fiebig & Lansner, 2017)?

The aim of the present study was to clarify the exact contribution of alpha oscillations to WM. In particular, we wanted to determine whether alpha band activity during WM maintenance primarily reflects attentional prioritization, or a more general maintenance mechanism. To address this question, we designed a visual WM experiment with functional impulse perturbation, in which we compared the maintenance of prioritized and deprioritized items. Priority was assigned by testing order, where the tested-first item was prioritized, while the tested-second item was deprioritized. We reasoned that if sustained alpha-band decoding reflects generic maintenance, it should consistently remain detectable for both items, even if its decoding strength differs to some extent with priority. Conversely, if it reflects attentional mechanisms, then we might still expect successful sustained decoding of tested-first items, but not of tested-second ones.

Our experiment featured two different retention intervals between the presentation of memory items and the subsequent memory probe: A short (1-second) and a long (3-second) interval. The rationale for incorporating an extended retention interval of up to 3 seconds was based on the idea of STSP as a maintenance mechanism in particular, as it predicts that residual calcium levels would progressively decline across an interval of up to one or two seconds (Mongillo et al., 2008; Zucker & Regehr, 2002), making longer intervals an ideal testcase for investigating potential neural reactivation mechanisms, which might be expressed in the alpha band specifically (Barbosa et al., 2021).

We were furthermore interested in the temporal dynamics of alpha oscillations during WM maintenance, as these might help to elucidate their functional role. Voltage-based decoding has previously shown that WM may maintain information through a stable coding scheme, while simultaneously exhibiting low-dimensional dynamic changes that reflect temporal progression or representational drift (Wolff, Jochim, et al., 2020). Assessing the temporal dynamics of alpha oscillations across both short and long delay intervals should provide information as to the nature and consistency of alpha oscillations during WM maintenance.

In addition to alpha band decoding, we decoded from the raw voltage signal, in line with prior impulse perturbation studies (Kandemir, Wilhelm, et al., 2024; Kandemir, Wolff, et al., 2024; Kandemir & Akyürek, 2023; Wolff et al., 2017; Wolff, Jochim, et al., 2020; Wolff, Kandemir, et al., 2020). Voltage decoding was meant to extend and replicate earlier reports of activity-quiescent maintenance and impulse-based decoding of memoranda, across both short and long delay intervals, providing a broadband reference for the alpha analyses.

## 2. Materials and methods

### 2.1 Participants

Thirty healthy adults were initially recruited for the electroencephalography (EEG) experiment. Two participants were excluded due to excessive noise in their EEG data (more than 30% of trials contaminated), leaving 28 participants (21 females, mean age 24 years, range 18-36 years). The study received ethical approval from the Ethical Committee of the Faculty of Behavioral and Social Sciences at the University of Groningen (Study ID = PSY-2122-S-0255). All participants gave written informed consent before participation and were compensated at a rate of €10 per hour.

### 2.2 Stimuli and apparatus

The experiment was conducted in a well-lit room, with the lighting set to a level of approximately 100 lux. The task was generated and presented using OpenSesame 3.3.14 (Mathôt et al., 2012), running on a standard Windows 10 desktop computer. Participants were seated about 60 cm away from a 27-inch ASUS VG279Q monitor, which had a refresh rate of 100 Hz and a resolution of 1920 by 1080 pixels. A USB keyboard was provided for the participants to enter their responses.

All stimuli were shown in the center of the screen, against a gray background (RGB = 128, 128,128). Among these was a black fixation dot, which spanned 0.25° of visual angle. Memory items and probes were sine-wave gratings presented at 20% contrast, with a diameter of 6.5° and a spatial frequency of 1 cycle per degree. Their phase was randomized within and across trials. The memory items had orientations of 0.5° to 178.0° in steps of 2.5°. The angle differences between memory items and the corresponding memory probes were uniformly distributed across 7 angle differences (±5°, ±10°, ±16°, ±24°, ±26°, ±32°, ±40°). The impulse stimuli consisted of white disks, with a diameter of 9.7°.

### 2.3 Experimental design and procedure

There were two blocked conditions in the experiment: The short maintenance delay condition (1 sec), and the long delay condition (3 sec). The experiment was divided into four consecutive sessions with a single day (18 blocks per session). At the beginning of each session, participants were informed which of the two memory items (the first- or second-presented item) would be probed first and which would be probed second. At the beginning of each block, a text (‘short’ or ‘long’) indicated the type of block to the participants.

The trial procedure is shown in Figure 1. Participants were instructed to maintain central fixation throughout the trials: each trial started with the presentation of the fixation dot, which remained on the screen for the entirety of the trial. After a 300-500 ms delay, two memory items were sequentially presented, each for 200 ms and separated by a 900 ms inter-stimulus interval. The maintenance delay followed (1 or 3 sec), after which the first impulse was presented for 100 ms. As in previous functional perturbation studies, the impulse serves to ‘re-illuminate’ WM representations during maintenance, even those that have become activity-quiescent during the preceding delay, thereby allowing us to decode their representations. Following another 500 ms delay, participants were probed on the “tested-first” memory item, with the probe being shown for 200 ms. The testing order was fixed within sessions 1-2 and within sessions 3-4, but switched between these two sets of sessions (i.e., the first-presented item was tested first in one pair of sessions, whereas the second-presented item was tested first in the other pair). This testing order was counterbalanced across participants. Participants indicated whether the orientation of probe relative to the memory item was rotated clockwise or counterclockwise by pressing the ‘m’ key, or the ‘c’ key, respectively. During the response period, participants were allotted a maximum of 1500 ms to respond. After a delay of 900 ms, the second impulse was presented for 100 ms. Finally, after another 500 ms delay, the second

**Fig 1.**
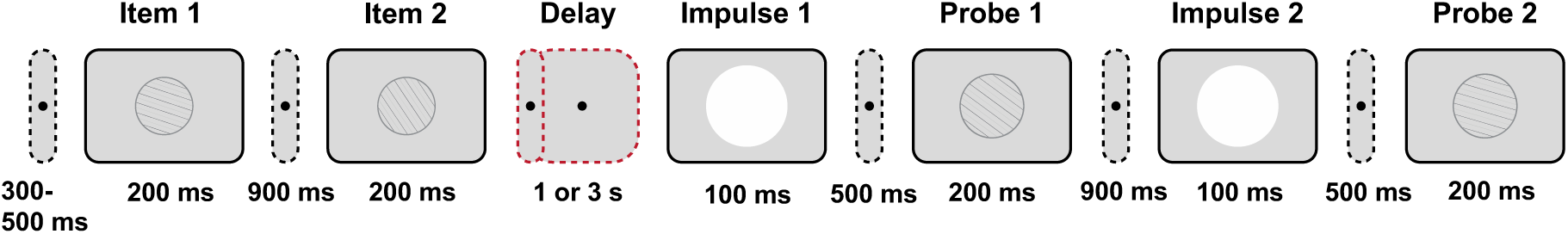
Trial schematic. Two randomly selected orientations were serially presented. Then two experimental conditions were implemented with either a short delay (1 s) or a long delay (3 s). After the delay, the first task-irrelevant impulse was presented, followed by a probe for the first (tested early) item. After an additional delay, the second task-irrelevant impulse was presented, followed by probing of the second (tested late) item.

memory probe was presented for 200 ms for the “tested-second” item. The next trial then followed automatically after the response. In addition to the main analyses based on test order (tested-first vs tested-second), we also report a control analysis, labeling items by presentation order (presented-first vs presented-second) in the Supplementary Materials.

In the EEG experiment, participants completed a total of 1728 trials, which were divided into 72 blocks and distributed across four consecutive sessions, all conducted within a single day. When participants completed a block of 24 trials, they were presented with feedback depicting their average accuracy for that block. Participants could take breaks between the blocks. The experiment had a total duration of approximately 6 hours, which included EEG cap preparation and participant-paced breaks.

### 2.4 EEG acquisition and preprocessing

The EEG data were acquired using 64 Ag/AgCl sintered electrodes, which were laid out according to the extended international 10-20 system. A TMSI Refa 8-64 amplifier was used to record EEG data at 1000 Hz, and CPz was used as online reference. Two additional electrodes were placed on the left and right mastoids, and the anterior midline frontal electrode (AFz) served as the ground. To record eye movements and blinks, the vertical electrooculogram (EOG) was derived from the voltage difference between an external electrode placed 2 cm below the left eye and the Fp1 electrode, while horizontal EOG was similarly obtained from the F9 and F10 electrodes. All electrode impedances were kept below 10 kΩ throughout the experiment.

The EEG data were pre-processed with EEGLAB v14.11 (Delorme & Makeig, 2004) and the FieldTrip toolbox (Oostenveld et al., 2011) in MATLAB. We first re-referenced the data to the average of right and left mastoids, downsampled to 500 Hz, and applied bandpass filtering (0.1-40 Hz). Then, the data were epoched relative to the onset of the first memory item (-200 ms to 900 ms), to the onset of the second memory item (short delay: -200 ms to 1200 ms, long delay: -200 ms to 3200 ms), and the onsets of the impulse stimuli (-200 ms to 600 ms). Each trial of each epoch was visually inspected for blinks, eye movements, and non-stereotypical artifacts, and epochs containing such artifacts were excluded from further analyses. Noisy channels were replaced through spherical spline interpolation. The average percentage of trials removed for each epoch were as follows: 10.77% in the item 1 epoch (short and long delay conditions together), 9.56% in the item 2 epoch (short and long delay conditions together), 9.88% in the short delay epoch, 12.91% in the long delay epoch, 7.54% in the impulse 1 epoch (short and long conditions together), and 7.65% in the impulse 2 epoch (short and long conditions together).

To extract alpha power amplitude, the EEG voltage data were first bandpass filtered between 8 to 12 Hz using the EEGLAB function eegfilt, which applies a two-pass (forward and backward), zero-phase finite impulse response (FIR) band-pass filter. The magnitude of the analytic signal was then extracted to obtain the amplitude envelope, constituting time-resolved alpha power. Induced alpha power was obtained by band-pass filtering single-trial data and applying a Hilbert transform before trial averaging, whereas evoked alpha power was computed using the same procedure after averaging voltage traces across trials. This was done separately for both induced and evoked alpha power. We focused primarily on induced alpha power because our hypotheses concern the oscillatory mechanism that sustain and prioritize items during the delay, rather than the stimulus-locked sensory response. Accordingly, we report induced alpha power in the main text, and the evoked alpha power analyses are provided in the Supplementary Materials. We computed induced alpha power by computing the alpha power of each trial’s voltage trace. Evoked alpha power reflects phase-locked neural responses, and was obtained by first averaging voltage traces across a subset of trials within each orientation bin (see decoding analyses below), followed by computing the alpha power on these averaged traces. To assess whether the outcomes were frequency-specific, we additionally repeated the same analyses in two neighboring bands: theta (4-7 Hz) and beta (14-30 Hz). These control analyses are reported in the Supplementary Materials.

### 2.5 Multivariate pattern analyses

We conducted several different multivariate pattern analyses (MVPA) on the raw voltage and alpha power signals to investigate the temporal dynamics of WM representations. Based on previous studies from our group, we selected 17 posterior channels (P7, P3, Pz, P4, P8, POz, O1, O2, P5, P1, P2, P6, PO3, PO4, PO7, PO8, Oz) for all decoding analyses (Kandemir, Wilhelm, et al., 2024; Kandemir & Akyürek, 2023; Wolff et al., 2017). These channels were chosen due to their relevance in capturing neural activity related to orientation processing in visual WM tasks. For reference, analyses were also conducted on data from all electrodes; these are reported in the Supplementary Materials. Decoding analyses were performed on all artifact-free trials and were not restricted to correct trials. Therefore, no trials were excluded based on behavioral accuracy.

The decoding analyses were performed to assess the neural representation of orientation information using different neural signals (time window of interest, raw voltage, and alpha power). Here, we present a concise summary of the shared methodological framework, followed by the specific differences characterizing each decoding analysis. The general decoding procedure involved baseline correction, down-sampling, orientation binning, cross-validated classification, and decoding accuracy computation. Firstly, baseline correction was performed by subtracting the mean signal computed within a pre-stimulus interval (-200 ms to 0 ms) from the entire time series for each trial and channel. To optimize computational efficiency, we then further downsampled the data to 100 Hz. For orientation decoding, trials were assigned to the closest one of 16 equal orientation angle bins by computing the minimal absolute distance between the stimulus orientation and the center values of the orientation bins. Trials were re-labeled with the center value of that bin. This binning process was repeated three times to cover the pre-determined orientation space (from 0° to 157.5°, 7.5° to 165°, and 15° to 172.5°, each in steps of 22.5°). This approach resulted in three runs, each with 16 different orientation conditions, allowing for a comprehensive assessment of orientation decoding across different bin alignments. These three runs were averaged to generate a final decoding result for each condition.

To decode the neural representation of orientation, we employed a cross-validated classification approach using Mahalanobis distance (De Maesschalck et al., 2000) as the metric. The trials were portioned into 8 folds using stratified K-fold cross-validation. In each iteration, seven folds served as the training set, and the remaining fold served as the test set. Within each training set, random subsampling was performed to equalize the number of trials across orientation conditions, and the data within each orientation condition were then averaged to create 16 condition-specific spatiotemporal patterns. To reduce noise and pool information across similar orientations, we convolved the spatiotemporal neural patterns associated with each orientation condition with a half-cosine basis set raised to the 15^th^ power (Kandemir, Wilhelm, et al., 2024; Kandemir, Wolff, et al., 2024; Wolff, Kandemir, et al., 2020). For each time point, pairwise Mahalanobis distances between the test trials and the averaged training patterns were computed. The covariance matrix estimated from the training data was computed by means of a shrinkage estimator (Ledoit & Wolf, 2003) to account for the variance structure of the data. Distances were mean-centered by subtracting the mean distance across all conditions, and sign-reversed, ensuring that positive values indicated higher similarity. For visualization, to summarize decoding accuracy, the Mahalanobis distances were sign-reversed and then averaged with weights based on the cosine of the orientation differences between orientations, spanning the full orientation space. The entire procedure was repeated 100 times with different random folds and subsampling to avoid sampling biases.

#### 2.5.1 Raw voltage time-course decoding

The raw voltage time-course decoding analysis aimed to capture the temporal dynamics of neural representations by decoding raw voltage data at each time point within the epoch. The specific decoding procedure was performed as described in section 2.5. The final decoding output was averaged across orientation spaces, repetitions and trials for each participant.

#### 2.5.2 Time window of interest decoding

To assess the overall decodability within specific epochs, channels and time samples within each predefined window were concatenated as features for decoding, and decoding performance was then averaged within the window. Specifically, we selected a time window of interest from 100 ms to 400 ms relative to the presentation of the relevant stimulus (i.e., memory item, or impulse). This time window was selected based on prior research indicating that the dynamic WM-dependent neural responses to visual stimuli occur mainly within this period (Wolff et al., 2017). Since we also aimed to assess WM maintenance, and to capture potential differences in sustained neural activity, we selected two further time-windows, corresponding to the end of the delay phase in each condition: 400-1000 ms for the short delay condition, and 2400-3000 ms for the long delay condition.

After adjusting the data as described in section 2.5, we reshaped it for the time window of interest analyses by concatenating the channel and time dimensions into one feature dimension. This enabled decoding of the interested time window as a unified pattern, rather than analyzing time point by time point. The reported decoding accuracy for each participant was calculated by averaging the weighted, mean-centered distances across all repetitions, trials and orientation spaces. This averaging yielded a single value per participant, representing the overall ability to decode orientation information from the neural data.

#### 2.5.3 Alpha power time-course decoding

The alpha power time-course decoding was highly similar to the decoding procedure of the raw voltage time-course decoding. For the alpha power time-course decoding, the analysis was conducted on alpha-band signals, separated into evoked and induced alpha power, following the same orientation binning procedure used in the preceding analyses. Then the data was divided into three folds using random stratified sampling, ensuring a balanced distribution of orientation bins across folds. To reduce trial-to-trial variability and enhance the reliability of decoding, pseudo-trials trials were generated by averaging trials within the same orientation bin for each fold. For induced alpha power decoding, trial-wise alpha power was computed before averaging, while for evoked alpha power decoding, voltage traces were averaged prior to filtering in the alpha frequency band.

For each time point, the classifier was trained on two folds (the training folds) and tested on the remaining fold (the test fold) using cross-validation. The covariance was computed from the training folds using a shrinkage estimator for stability. In the training folds, averaged pseudo-trials for each orientation bin were smoothed using a half-cosine basis function to pool neighboring orientations, raising it to the 15^th^ power to enhance selectivity. The Mahalanobis distances were computed between each test pseudo-trial and the averaged training pseudo-trials across orientation bins. These distances were cosine-weighted, sign-reversed, and mean-centered to generate a similarity profile. A cosine vector mean was computed to summarize the similarity, where higher values indicated greater similarity between the neural representations of the tested and trained orientations, reflecting successful decoding of the orientation information from the alpha power signal. The entire decoding process was repeated 100 times with randomized fold assignments to ensure robustness. Decoding accuracy was also averaged over iterations, orientation spaces, and time points for each participant. The final decoding accuracy values for evoked and induced alpha power were averaged across all iterations, providing a stable estimate of the temporal dynamics of the alpha power representation for each participant.

#### 2.5.4 Cross-temporal generalization

To further investigate the temporal dynamics of WM content, we conducted cross-temporal generalization analyses (King & Dehaene, 2014), focusing specifically on decoding performance for the tested-first item during the delay period. The decoding methodology remained consistent with the procedures outlined previously, except for the classifier’s training and testing regimen. Specifically, instead of training and testing the classifier at each corresponding time point, we trained the classifier at a specific time point and then tested it across all other time points. This resulted in a two-dimensional temporal generalization matrix that provides classification accuracy for every combination of training and testing times, indicating whether neural coding is dynamic or static over time.

### 2.6 Statistical assessment

We applied repeated-measure ANOVAs and made planned comparisons between the delay conditions for tested-first and tested-second items with paired samples t-tests, to test the differences in behavioral performance. To evaluate the significance of our decoding results across the different analyses, we employed permutation tests to generate null distributions against which the observed decoding performance was compared. Specifically, we performed 100,000 permutations where the sign of the decoding accuracy for each participant was randomly flipped with 50% probability. The proportion of the null distribution where the permuted mean decoding accuracy was greater than the observed mean decoding accuracy was calculated as the *p*-value. The decoding performance was considered statistically significant if this *p*-value was less than 0.05.

For the time-course decoding analysis and time window of interest decoding, a cluster-based correction was applied to control for multiple comparisons, identifying clusters of consecutive time points where decoding performance exceeded chance levels. Bootstrapped confidence intervals were computed to estimate the variability of the mean decoding performance over time. All tests of the time-course decoding and time window of interest decoding tests were one-sided, as negative values are not meaningful. We again used paired samples t-tests to test differences between the short and long delay conditions, as well as differences between tested-first and tested-second items within each condition.

To quantify evidence for above-chance decoding in the predefined late delay window (0.4-1 s or 2.4 - 3.0 s), Bayes factors (BF) were computed, using the freely available JASP software (JASP Team, 2024) We conducted Bayesian one-sample t-tests against 0, where decoding values were converted to deviations from chance by subtracting chance level. Bayes factors are reported as BF_10_.

For the cross-temporal generalization analysis, a cluster-based permutation test was employed as well. Difference matrices were computed by subtracting diagonal decoding accuracies from off-diagonal elements along both training and testing time dimensions, to assess the temporal generalization of neural representations.

## 3. Results

### 3.1 Behavioral results

A 2 × 2 repeated-measure ANOVA was conducted to examine the effects of test order (tested-first vs. tested-second) and retention interval (1s short-delay vs. 3s long-delay) on response accuracy (Figure 2). The analysis revealed that accuracy was significantly higher for the tested-first item than the tested-second item, *F* (1, 27) = 207.67, *p* < 0.001, η²p = 0.885. However, the retention interval had no significant effect on accuracy, *F* (1, 27) = 1.44, *p* = 0.240, η²p = 0.051. Importantly, there was a significant interaction between test order and retention interval, *F* (1, 27) = 13.02, *p* = 0.001, η²p = 0.325, suggesting that the effect of delay differed between the tested-first and tested-second items.

**Figure 2.**
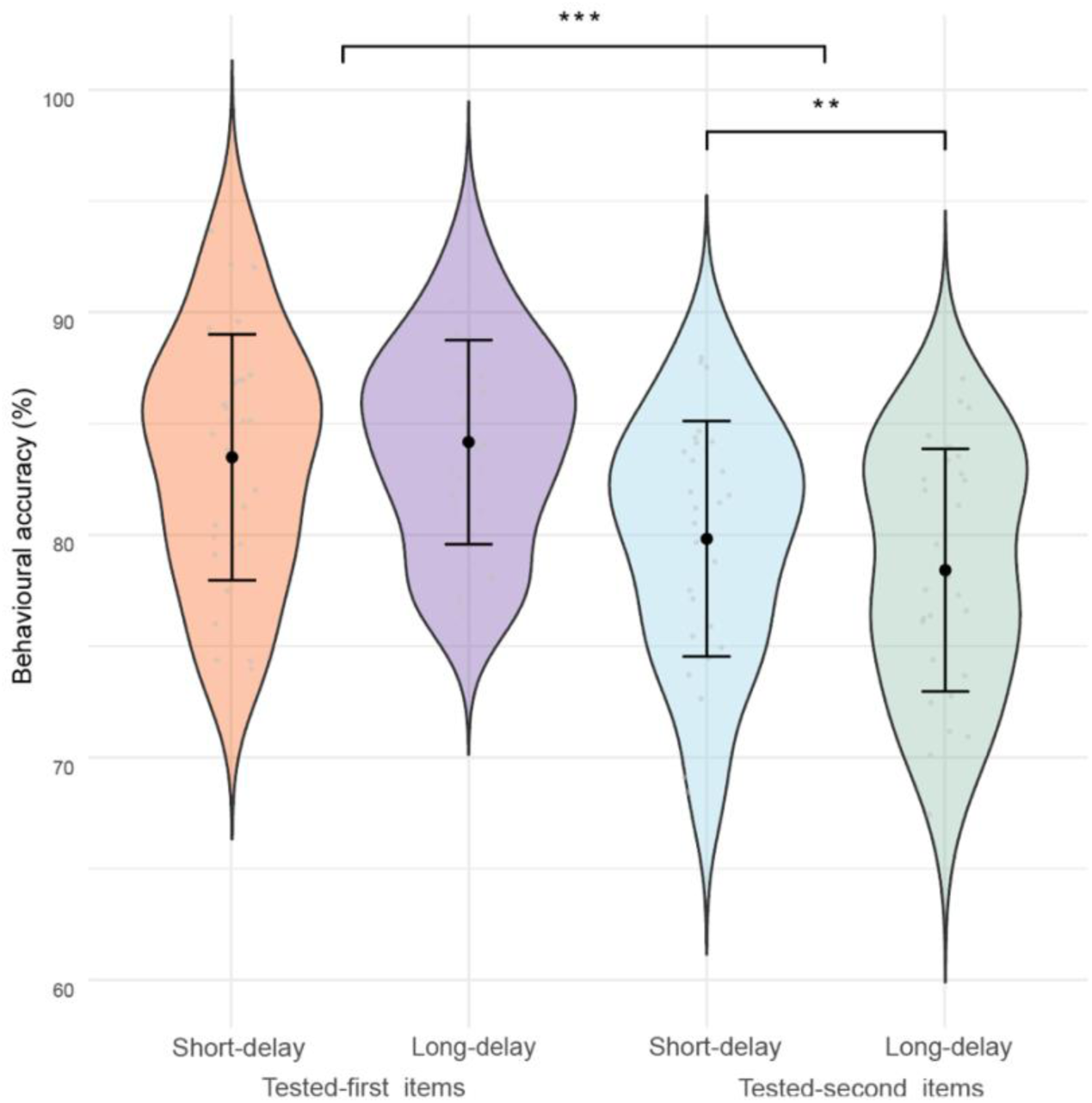
Violin plot representing response accuracy (%) for the tested-first and tested-second items in the short (1s) and long delay (3s) conditions. The width of each violin indicates the distribution density of accuracy values across participants. The black dots represent the mean accuracy, and the error bars indicate the 95% confidence intervals. Asterisks indicate statistically significant effects (***, *p* < 0.001; **, *p* < 0.01).

To unpack this interaction, we conducted follow-up paired-samples t-tests. For the tested-first item, accuracy did not differ between the long-delay (*M* = 0.8427, *SD* = 0.0480) and short-delay conditions (*M* = 0.8371, *SD* = 0.0559, *t* (27) = -1.39, *p* = 0.175, *Cohen’s d* = - 0.26). However, a significant effect was observed on the tested-second item: accuracy in the short-delay condition (*M* = 0.8002, *SD* = 0.0491) was significantly higher than in the long-delay condition (*M* = 0.7879, *SD* = 0.0545), *t (27)* = 3.61, *p* = 0.001, *Cohen’s d* = 0.68). These findings suggest that while the duration of the delay period did not influence retrieval performance for the tested-first item, it had a measurable impact on the tested-second item, with prolonged delays leading to slightly reduced accuracy (from 80.02% to 78.79%).

### 3.2 Voltage decoding of memory encoding

We first performed a time course decoding analysis to test whether our participants differentially encoded the memory items depending on the delay period they were expecting. Across both short and long delay conditions, as expected, the representations of item 1 and item 2 exhibited clear parametric similarity patterns, as illustrated in Figure 3A and Figure 3D. In the short delay condition, as shown in Figure 3A-C, item 1 could be decoded (Figure 3B, left, 62-1100 ms, *p* < 0.001, and Figure 3C, left, *p* < 0.001, both one-tailed, corrected) as well as item 2 after its respective presentation (Figure 3B, right, 72-1100 ms, *p* < 0.001, and Figure 3C, right, *p* < 0.001, both one-tailed, corrected).

**Figure 3.**
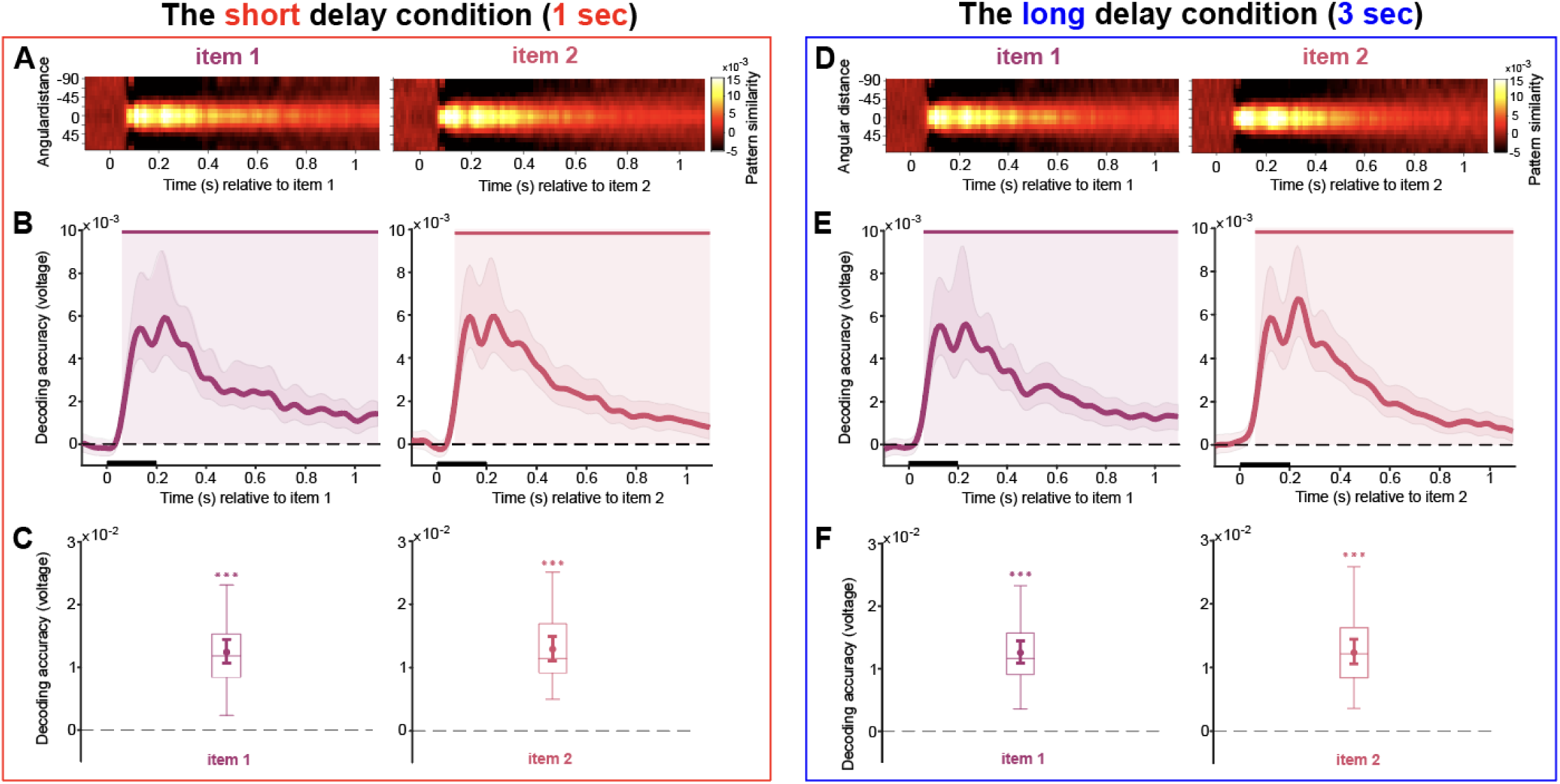
Decoding during item encoding. A-C, the short delay condition. D-F, the long delay condition. A and D, pattern similarity was quantified by computing mean-centered, sign-reversed Mahalanobis distances, averaged across trials for each time point, yielding a normalized measure of orientation-specific neural representations. B and E, decoding accuracy of memory item 1 and 2 during memory item presentation. Upper solid bars and corresponding shading indicate significant decoding areas (one-sided, *p* < 0.05). Error shading indicates 95% CI of the mean. C and F, boxplots indicating decoding accuracy for memory items within the 100-400 ms time window of interest relative to the onset of item 1 and 2. The middle line within each box represents the median, while the box itself spans the 25^th^ to 75^th^ percentiles. Whiskers extend to 1.5 times the interquartile range. Mean decoding accuracy is marked by filled circles, with vertical error bars showing the 95% CI. Asterisks indicate decoding accuracies that are significantly above zero (***, *p* < 0.001).

In the long delay condition, shown in Figure 3D-F, decoding performance was very similar, and we observed robust decoding of memory item 1 (Figure 3E, left, 60-1100 ms, *p* < 0.001, and Figure 3F, left, *p* < 0.001, both one-tailed, corrected) and item 2 (Figure 3E, right, 63-1100 ms, *p* < 0.001, and Figure 3F, right, *p* < 0.001, both one-tailed, corrected).

Comparing overall decoding in the time window of interest between the short delay and long delay conditions, decoding of item 2 appeared to be slightly stronger in the short delay condition (*t* (27) = 2.10, *p* = 0.045, *Cohen’s d* = 0.35), whereas no difference was observed for item 1 (*p* > 0.05).

In summary, these results reveal a robust and sustained neural representation during memory encoding of both item 1 and item 2 in the short and long delay conditions, with decoding performance remaining largely consistent, even though there was a slight enhancement for item 2 in the short delay condition. This indicates a consistent encoding of orientation-specific information across varying delay durations.

### 3.3 Delay interval decoding

#### 3.3.1 Voltage

While in the above we examined memory encoding by directly assessing the representations of the memory items in the order of their *presentation*, to assess maintenance of information in WM, the analyses below are based on the (session-wise) *report order* of the items, as the latter should reflect the priority of the items in WM. These analyses started after the presentation of item 2, when both items were available to the participants, and assessed their representations across the length of the respective delay intervals.

In the short delay condition, consistent with previous studies (Wolff et al., 2017; Wolff, Kandemir, et al., 2020), decoding accuracy of both the tested-first (Figure 4A, left, 166-740 ms, and 778-843 ms, *p* < 0.001, one-sided, corrected) and tested-second item (Figure 4A, left, 166-694 ms, *p* < 0.001, one-sided, corrected) exhibited an early peak shortly after the onset of the second memory item, with the tested-first item having a smaller significant cluster towards the end of the delay interval as well. In the analysis of our predefined time windows of interest, both the tested-first and tested-second items were decodable in the early (100-400 ms), as well as in the late (400-1000 ms) time window (Figure 4A, right, all *p* < 0.001, one-sided; late-window: tested-first: BF_10_ = 19.114, tested-second: BF_10_ = 2.847).

**Figure 4.**
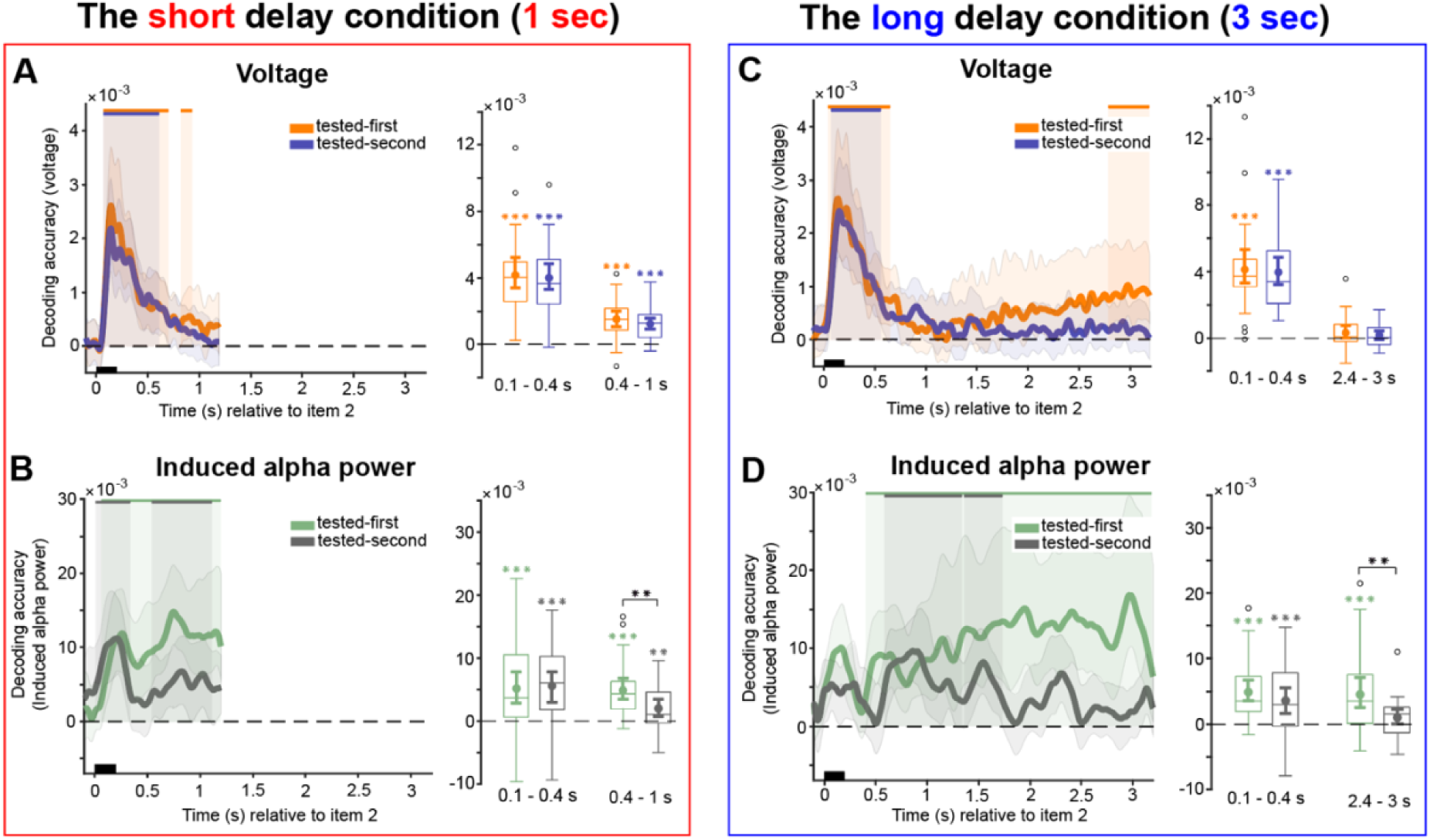
Decoding during delay intervals. A-B, the short delay condition. C-D, the long delay condition. A and C, raw voltage time course decoding. B and D, induced alpha power time course decoding. Upper solid bars and corresponding shading indicate significant decoding areas (one-sided, *p* < 0.05). Error shading represents 95% CI of the mean. Boxplots illustrate decoding accuracy for tested-first and tested-second items across the two selected time windows of interest in the short delay condition (100-400 ms and 400-1000 ms) and the long delay condition (100-400 ms and 2400-3000 ms). The middle line within each box represents the median, while the box itself spans the 25^th^ to 75^th^ percentiles. Whiskers extend to 1.5 times the interquartile range, and individual dots indicate extreme values. Mean decoding accuracy is marked by filled circles, with vertical error bars showing the 95% CI. Asterisks indicate significant decoding performance (*, *p* < 0.05; **, *p* < 0.01; ***, *p* < 0.001).

In the long delay condition, both the tested-first (Figure 4D, left, 28-693 ms, 2783-3200ms, *p* < 0.001, one-sided, corrected) and tested-second items (Figure 4D, left, 43-564 ms, *p* < 0.001, one-sided, corrected) were decodable immediately after the onset of the memory item 2, with the tested-first item again showing a slightly later offset. However, decoding of the tested-first item seemed to ramp up towards the end of the interval, presumably in preparation of the moment when the probe would appear, and critically, the tested-first item could be decoded again at that moment, while the tested-second item could not. The analysis of the early time window of interest (100-400 ms) revealed that both tested-first and tested-second items were decodable (Figure 4D, right, both *p* < 0.001, one-sided). In the late time window (2400-3000 ms), the tested-first item exhibited a marginal effect (Figure 4D, right, *p* = 0.05, one-sided, BF_10_ = 3.137), while the tested-second item did not reach significance (Figure 4D, right, both *p* = 0.108, one-sided, BF_10_ = 0.505).

To determine whether decoding accuracy differed between tested-first and tested-second items, we conducted further pairwise t-tests within both delay conditions. No significant differences were found between tested-first and tested-second items in the short or long delay conditions for any of the predefined time windows (all *p* > 0.05). Given the potential impact of the duration of the maintenance interval on neural representations, we also examined whether decoding accuracy differed between short and long delay conditions. We again observed no significant differences between short and long delay conditions for tested-first and tested-second items within the early time window (all *p* > 0.05). However, in the late time window, decoding accuracy for both tested-first (*t* (27) = 3.52, *p* < 0.01, *Cohen’s d* = 0.65) and tested-second items (*t* (27) = 5.05, *p* < 0.001, *Cohen’s d* = 0.93) in the short delay condition was significantly higher than in the long delay condition.

Overall, the raw voltage-based analyses largely replicated previous studies that have observed limited to no decoding after the initial stimulus-evoked response (e.g., Wolff et al., 2017; 2020a; 2020b). In the long delay condition in particular, decoding accuracy of the tested-first item showed a steady increase towards the end of the interval, presumably in anticipation of the ensuing probe. In line with this interpretation, no such increase was observed for the tested-second item, for which the response was not yet due at that point.

#### 3.3.2 Alpha power

Induced alpha power decoding in the short delay interval revealed that the tested-first item exhibited continuous decodability, whereas the tested-second item showed two shorter stretches of decodability (Figure 4C, left; tested-first: 66-1198 ms, *p* < 0.001; tested-second: 10-336 ms, *p* = 0.004, and 536-1108 ms, *p* = 0.003, both one-sided, corrected). However, our time-window analyses did not further support these differences: Tested-first as well tested-second items could be decoded in both the early and late time windows (Figure 4C, right; 100-400 ms: tested-first: *p* < 0.001; tested-second: *p* < 0.001; 400-1000 ms: tested-first: *p* < 0.001, BF_10_ > 1000; tested-second: *p* = 0.003; all one-sided, BF_10_ = 8.314).

Induced alpha power in the long-delay condition showed that the tested-first item was decodable during a substantial portion of the delay interval (Figure 4F, left, 406-3182 ms, *p* < 0.001, one-sided, corrected). By contrast, decoding of the tested-second item was more transient, with significant clusters early in the delay (Figure 4F, left, 584-1340 ms, *p* < 0.001, and 1362-1738 ms, *p* = 0.018, both one-sided, corrected). Across the early time window of interest, both tested-first (Figure 4F, right, *p* < 0.001, one-sided) and tested-second items (Figure 4F, right, *p* < 0.01, one-sided) could nevertheless be decoded. However, in the late time window, decoding remained reliable for the tested-first item (Figure 4F, right, *p* < 0.01, one-sided, BF_10_ = 21.219), while the tested-second item did not show reliable above-chance decoding (Figure 4F, right, *p* > 0.05, one-sided, BF_10_ = 0.586).

A direct comparison of tested-first and tested-second items revealed a significant difference in the late time window of interest for the both short and long delay intervals, with lower induced alpha power decoding accuracy for the tested-second item (short delay: *t* (27) = 3.03, *p* < 0.01, *Cohen’s d* = 0.57; long delay: *t* (27) = 2.99, *p* < 0.01, *Cohen’s d* = 0.56). No differences were observed in other cases (all *p* > 0.05). We further examined whether decoding accuracy differed between short and long delay conditions. No differences in induced alpha power between the short and long delay conditions, in either time windows, reached significance (all *p* > 0.10).

Taken together, the analyses of induced alpha power are compatible with the idea that internally generated alpha waves play a role in the sustained, attentional maintenance of prioritized (tested-first) items in WM specifically. As a general mechanism to maintain information in WM (i.e., pertaining to both prioritized and deprioritized items), it would nevertheless seem to fall short, as we did not find sustained alpha oscillations related to deprioritized (tested-second) items.

### 3.4 Impulse 1

#### 3.4.1 Voltage

Decoding voltage in the impulse 1 epoch in the short delay condition revealed that the tested-first item remained decodable throughout the post-impulse period (Figure 5A, left, 72-600 ms, *p* < 0.001, one-tailed, corrected), while the tested-second item, which had become completely undecodable prior to impulse 1 onset, also exhibited a brief period of significant decoding, peaking at approximately 200 ms post-impulse onset, and quickly returning to chance level (Figure 5A, left, 182-242 ms, *p* = 0.006, one-tailed, corrected). The analysis of the time window of interest (100-400 ms relative to impulse 1 onset) showed similar results. Both the tested-first (Figure 5A, right, *p* < 0.001, one-sided) and tested-second items (Figure 5A, right, *p* = 0.023, one-sided) were decodable, yet the tested-first item exhibited a stronger representational signal compared to the tested-second item (*t* (27) = 5.07, *p* < 0.001, *Cohen’s d* = 1.11).

**Figure 5.**
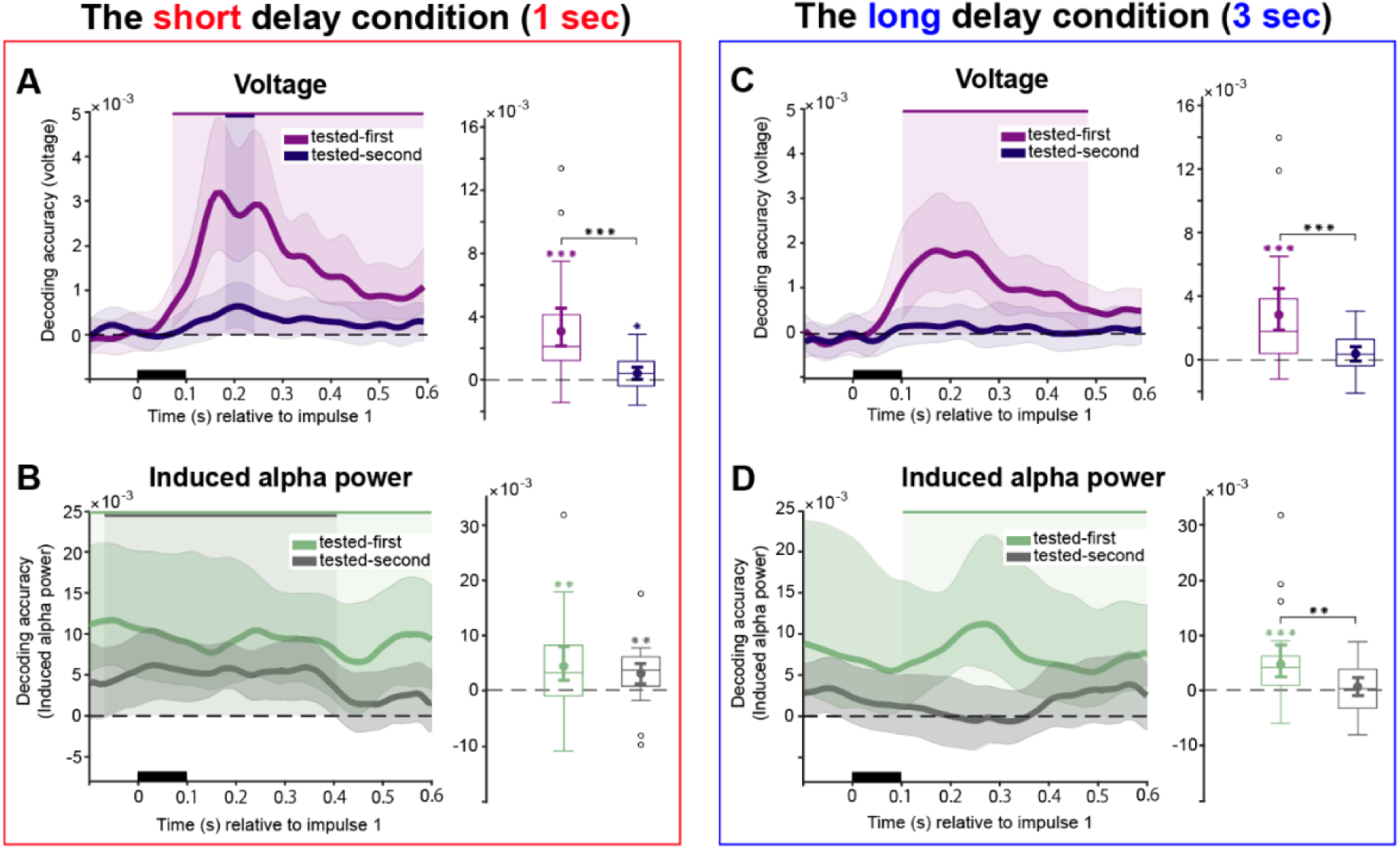
Decoding content from the first impulse response. A-B, the short delay condition. C-D, the long delay condition. A and C, raw voltage time course decoding. B and D, induced alpha power time course decoding. Upper solid bars and corresponding shading indicate significant decoding areas (one-sided, *p* < 0.05). Error shading represents 95% CI of the mean. Boxplots illustrate decoding accuracy for tested-first and tested-second items within the time window of interest (100-400 ms). The middle line within each box represents the median, while the box itself spans the 25^th^ to 75^th^ percentiles. Whiskers extend to 1.5 times the interquartile range, and individual dots indicate extreme values. Mean decoding accuracy is marked by filled circles, with vertical error bars showing the 95% CI. Asterisks indicate significant decoding performance (*, *p* < 0.05; **, *p* < 0.01; ***, *p* < 0.001).

In the long delay interval, raw voltage time course decoding showed a significant response for the tested-first item (Figure 5D, left, 102-482 ms, *p* < 0.001, one-tailed, corrected), but not for the tested-second item, even though participants responded accurately to the probe afterwards. The time-of-interest (100-400 ms) analysis also revealed that only the tested-first item could be decoded (Figure 5D, right, *p* < 0.001, one-sided), while the tested-second item could not (Figure 5D, right, *p* = 0.060, one-sided). Moreover, a clear difference in decoding strength was evident between the two items (*t* (27) = 4.29, *p* < 0.001, *Cohen’s d* = 0.81).

#### 3.4.2 Alpha power

In the short delay condition, the decoding results for induced alpha power showed that both the tested-first item (Figure 5C, left, -100-600 ms, *p* < 0.001, one-sided, corrected) and the tested-second item (Figure 5C, left, -68-406 ms, *p* < 0.001, one-sided, corrected) were decodable. Within the time window of interest, both the tested-first (Figure 5C, right, *p* < 0.01, one-sided) and tested-second items (Figure 6C, right, *p* < 0.01, one-sided) were decodable (*t* (27) = 0.76, *p* = 0.454, *Cohen’s d* = 0.14).

**Figure 6.**
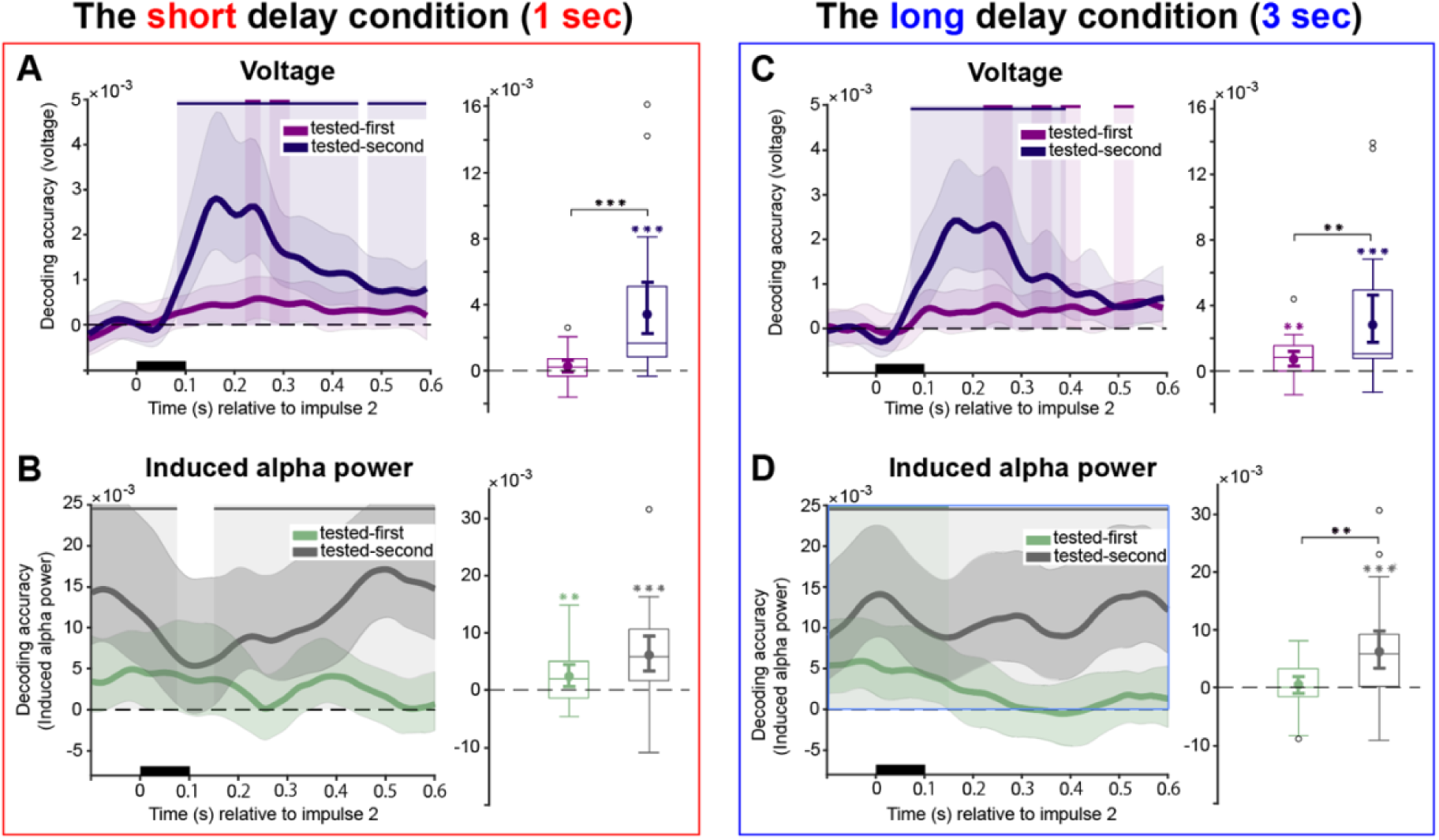
Decoding content from the second impulse response. A-B, the short delay condition. C-D, the long delay condition. A and C, raw voltage time course decoding. B and D, induced alpha power time course decoding. Upper solid bars and corresponding shading indicate significant decoding areas (one-sided, *p* < 0.05). Error shading represents 95% CI of the mean. Boxplots illustrate decoding accuracy for tested-first and tested-second items within the time window of interest (100-400 ms). The middle line within each box represents the median, while the box itself spans the 25^th^ to 75^th^ percentiles. Whiskers extend to 1.5 times the interquartile range, and individual dots indicate extreme values. Mean decoding accuracy is marked by filled circles, with vertical error bars showing the 95% CI. Asterisks indicate significant decoding performance (*, *p* < 0.05; **, *p* < 0.01; ***, *p* < 0.001).

In the long delay condition, induced alpha power time-course analysis revealed that only the tested-first item was decodable from induced alpha power following impulse 1 (Figure 5F, left, 104-600 ms, *p* = 0.001, one-sided, corrected). Across the 100-400 ms time window of interest, significant decoding emerged for the tested-first item (Figure 5F, right, *p* < 0.001, one-sided), as well as a significant difference between tested-first and tested-second items (*t* (27) = 3.36, *p* = 0.002, *Cohen’s d* = 0.64).

### 3.5 Impulse 2

#### 3.5.1 Voltage

The raw voltage time course decoding analysis for impulse 2 in the short delay condition revealed that the tested-second item was decodable throughout almost the entire delay period (Figure 6A, right, 82-452 ms, *p* < 0.001, and 472-600 ms, *p* = 0.032, one-tailed, corrected). Note that this pattern for the tested-second item is not unexpected, as at this stage, this item had become task-relevant for the upcoming second probe. By contrast, the tested-first item was only decodable within two narrow time windows (Figure 6A, right, 222-252 ms, *p* = 0.037, and 272-312 ms, *p* = 0.033, one-tailed, corrected). The analysis of the time window of interest (100-400 ms relative to impulse 2 onset) demonstrated that only the tested-second item could be successfully decoded (Figure 6A, right, *p* < 0.001, one-sided), whereas the tested-first item could not (Figure 6A, right, *p* = 0.080, one-sided). Moreover, there was a marked difference in decoding strength between the two items (*t* (27) = -4.06, *p* < 0.001, *Cohen’s d* = 0.76).

Decoding results for impulse 2 in the long delay showed that the tested-second item was consistently decodable for about 300 ms after impulse 2 onset (Figure 6D, right, 72-392 ms, *p* < 0.001, one-tailed, corrected), while the tested-first item was again only significantly decoded in short, discrete time windows (Figure 6D, right, 222-282 ms, *p* = 0.01, 322-362 ms, *p* = 0.047, 382-422 ms, *p* = 0.03, and 492-532 ms, *p* = 0.025, one-tailed, corrected). In the time window of interest (100-400 ms) both the tested-first item (Figure 6D, right, *p* < 0.001, one-sided) and tested-second items were decodable (Figure 6D, right, *p* = 0.060, one-sided), although the tested-second item exhibited a higher decoding strength (*t* (27) = 3.24, *p* = 0.003, *Cohen’s d* = 0.82).

#### 3.5.2 Alpha power

In the short delay condition, the tested-second item could be decoded from induced alpha power (Figure 6C, left, -100-74 ms, *p* = 0.035, and 150-600 ms, *p* < 0.001, one-sided, corrected), with decoding accuracy remaining above chance for a sustained period after impulse 2 presentation. In the time window of interest, both the tested-first (Figure 6C, right, *p* = 0.01, one-sided) and tested-second items (Figure 6C, right, *p* < 0.001, one-sided) remained decodable from induced alpha power, without a significant difference between them (*t* (27) = -1.86, *p* = 0.074, *Cohen’s d* = -0.36).

A similar pattern emerged in the long delay condition, with reliable decoding of the tested-second item from induced alpha power (Figure 6F, left, -100-600 ms, *p* < 0.001, one-sided, corrected). In addition, the tested-first item was decodable from induced alpha power during an early, short interval (Figure 6F, left, -100-148 ms, *p* = 0.005, one-sided, corrected). Decoding within the predefined time widow also indicated significant decoding for the tested-second item only, from induced alpha power (Figure 6F, right, *p* < 0.001, one-sided). Importantly, there was again a significant difference between tested-first and tested-second items in induced alpha power (*t* (27) = -2.88, *p* = 0.008, *Cohen’s d* = -0.54). No further significant group-level differences were detected between the short and long delay conditions for either the tested-first or tested-second items, across all signals (voltage and induced alpha power) and across both impulses.

### 3.6 Cross-temporal generalization

#### 3.6.1 Voltage

The results revealed striking differences in temporal stability between the short and long delay conditions when analyzing raw voltage data. The short delay epoch exhibited highly dynamic decoding patterns where tested-first item representations changed rapidly over time. In the cross-temporal matrix, above-chance effects were concentrated along the diagonal, while generalization away from the diagonal remained sparse and spatially limited. Consistent with this pattern, decoding at off-diagonal time points was reliably lower than decoding on the diagonal, supporting a highly dynamic representational format during the short delay interval (Figure 7A, cluster-based permutation test, cluster-corrected, *p* < 0.05). The long-delay condition elicited an initial dynamic state similar to that in the short delay. From approximately midway through the delay period, decoding shifted toward a more stable representational pattern, expressed as a broad region of sustained above-chance decoding that extended beyond the diagonal and was absent in the short delay condition. When decoding away from the diagonal was compared with same time decoding, off-diagonal performance was relatively less reduced during this later period, consistent with stronger cross-temporal generalization than in the short delay condition (Figure 7D, cluster-based permutation test, cluster-corrected, *p* < 0.05).

**Figure 7.**
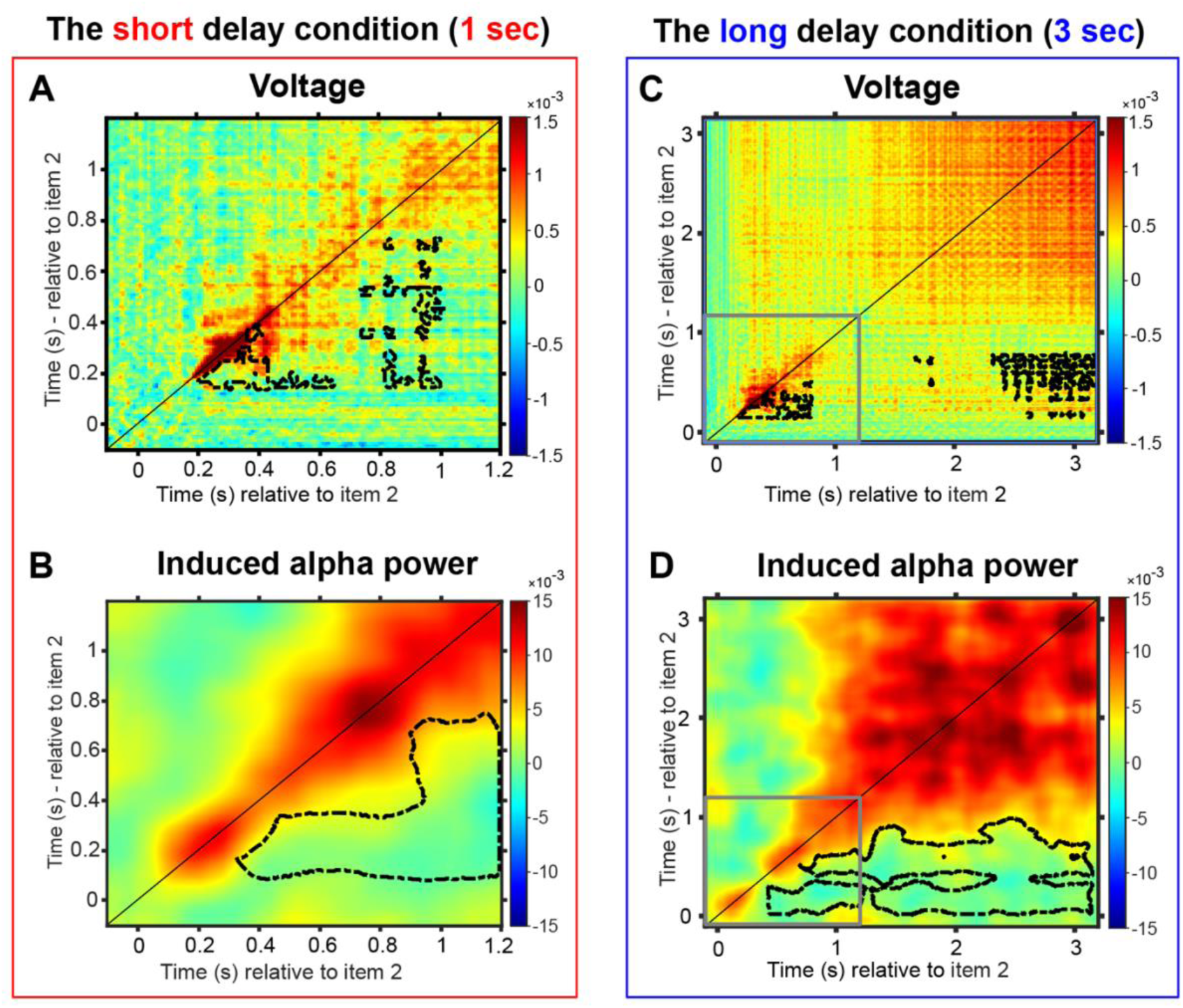
Cross-temporal decoding matrices for the tested-first item, derived from training and testing on all time-point combinations. A-B, the short delay condition. C-D, the long delay condition. A and C, cross-temporal generalization in raw voltage. B and D, cross-temporal generalization in induced alpha power. Decoding accuracy is represented by color intensity. The black lines indicate time-points of significantly lower decoding relative to both equivalent time-points along the diagonal (same train and test time point, *p* < 0.05, cluster corrected). Note the scale difference illustrated by the grey outline in panels C and D, which corresponds to the extent of time plotted in panels A and B.

#### 3.6.2 Alpha power

In the short delay condition, induced alpha decoding exceeded chance in a single extended region that emerged after approximately 300 ms and largely followed the diagonal, with only a narrow spread to adjacent off-diagonal time points. This limited cross-temporal transfer suggests that the underlying representational format was dynamic, rapidly changing over time. When comparing decoding away from the diagonal to same-time decoding, off-diagonal decoding was reliably reduced, consistent with a largely time-specific code (Figure 7C, cluster-based permutation test, cluster-corrected, *p* < 0.05). In the long delay condition, while the initial dynamic, diagonal-only pattern was also observed in the early stage, a clear shift toward a more stable neural representation emerged approximately one second post-stimulus. Consistent with this transition, above-chance decoding extended beyond the diagonal across a broad portion of the train-by-test matrix, indicating increased cross-temporal generalization during the delay. When decoding away from the diagonal was compared to same time decoding, off-diagonal decoding was less attenuated in the long delay condition, further supporting a more stable representational format over time (Figure 7F, cluster-based permutation test, cluster-corrected, *p* < 0.05). This finding suggests a fundamental change in how WM representations are coded across time: In the short delay condition, neural representations remain transient and dynamic, whereas in the long delay condition, they gradually stabilize, potentially reflecting a distinct neural mechanism for prolonged maintenance.

## 4. Discussion

To investigate how alpha-band activity supports the maintenance of WM items, we compared maintenance of prioritized (tested-first) and deprioritized (tested-second) items, across different delays between memory encoding and retrieval, comparing short (1-second) and long (3-second) retention intervals. This design allowed us to test whether alpha oscillations primarily index selective attentional prioritization in WM^2^, or rather support a broader maintenance process, to keep memoranda persistently activated, or at least periodically refreshed. Our analyses demonstrated a clear dissociation between behaviorally tested-first and tested-second memory items, supporting the former account.

Our analyses of the long delay interval in particular showed how priority was supported by alpha oscillations. Specifically, while orientation-specific information for both tested-first and tested-second items was initially decodable after encoding, only the tested-first item remained decodable throughout the extended three-second interval, from induced alpha power exclusively. By contrast, the tested-second item showed no robust evidence of sustained decodability in the late delay, either in raw voltage or in alpha band power, which is consistent with a transition toward a less persistent representational state. That said, in the first half of the long delay interval, the tested-second item was also visible in induced alpha, potentially indicating residual attentional maintenance of this item. While less univocal in some respects, the patterns observed in the short delay interval condition were qualitatively similar. The overall priority-based difference between the items during the maintenance delay suggests that sustained alpha oscillations mainly support attentional prioritization rather than general maintenance in WM.

Impulse response decoding also closely tracked priority; decoding of the tested-first item was strong in all measures from the first impulse, and while the tested-second item could also be detected, it was clearly much weaker. At the second impulse the picture was reversed, where decoding of the now-relevant tested-second item was strong, and that of the obsoleted tested-first item was very weak to absent entirely. Thus, when the tested-second item became task-relevant again to select a response to the second probe, and even though it had been absent prior, it could be decoded from the preceding, second impulse.

Our findings align with broader theoretical frameworks proposing that alpha oscillations are involved in top-down attentional control and sensory gating (Bonnefond & Jensen, 2012; Jensen & Mazaheri, 2010; Klimesch, 2012). Accordingly, in the context of WM, alpha band oscillations are thought to play a role in guiding goal-directed action (Myers et al., 2017; van Ede et al., 2017), and have also been associated with inhibition of irrelevant information and/or further visual input in order to protect memory contents (Payne & Sekuler, 2014; Sauseng et al., 2009; Wianda & Ross, 2019). Although the idea that attention might support WM maintenance in this manner has intuitive appeal, it bears noting that the present results suggest that these functions are exclusively reserved for items that are prioritized in WM, not all items. In other words, the boundaries of attention are not the boundaries of WM (cf. Oberauer & Awh, 2022), which is contrary to some previous proposals (Chun, 2011; Cowan, 2001; Foster & Awh, 2019). The selective role of attention nevertheless fits well with proposals of prioritized (or “activated”) WM, which is thought to constitute a privileged subset of WM (Borst et al., 2010; Manohar et al., 2019; Oberauer, 2002; Olivers et al., 2011), and which may exist without the need to sacrifice retention of other memoranda in WM (Myers et al., 2018).

As indicated, some previous studies have suggested a broader, more general role for alpha oscillations in WM maintenance. Foster and colleagues (2017; Sutterer et al., 2019) reported persistently active spatial representations in the alpha band, even arising regardless of task relevance, which suggest these oscillations might serve a more general maintenance process. Ten Oever and colleagues (2020) found that alpha phase modulated the EEG impulse response during WM maintenance, attributing the effect to an interaction with ongoing activity. Similarly, Barbosa and colleagues (2021) decoded WM items from alpha band power, and interpreted this as a sign of representational maintenance based on ongoing activity. Chen and colleagues (2023) even reported that non-invasive modulation of parietal alpha oscillations during WM retention can alter WM performance, consistent with a causal contribution of parietal alpha to WM retention. However, despite these prior findings, the idea that persistent alpha oscillations support WM maintenance in general, either as an active representational code, or as a refreshing mechanism to keep synaptic representations in place, was generally not supported by our data, especially when considering the decoding patterns in the long delay interval.

We can think of three main reasons for why our results might diverge from those of these previous studies that seemed to suggest a more general role for alpha oscillations in WM maintenance. First, we purposefully implemented a longer delay condition to tap into processes that might play out beyond the earliest phase of WM maintenance, during the first one or two seconds. We reasoned that this early phase might be special, for instance because calcium-mediated short-term synaptic connectivity would be effective across this interval, but not beyond, as the supporting calcium residuals would then have dissipated (Mongillo et al., 2008; Pals et al., 2020). Previous analyses of WM maintenance have typically used short delay intervals within this relatively short range (e.g., Kandemir, Wilhelm, et al., 2024; Yang et al., 2025), which might have produced an incomplete picture of the maintenance process. In the present data, the outcomes in the short interval were also more ambiguous with regard to sustained activity, where we observed successful, albeit intermittent, decoding of the tested-second item from alpha band power, for instance.

Second, studies of WM have often relied on tasks in which only a single item is retained during the delay interval (e.g., Thrower et al., 2023), which also applies to tasks in which one item is singled out from a larger set for eventual recall by a cue, and in which the other items can then be discarded altogether (e.g., F.-W. Chen et al., 2023; Karabay et al., 2025). When there is only a single task-relevant item, and particularly when attention has been called towards that item by a cue stimulus, it seems likely that it will stay in the focus of attention. In the present study, we were able to contrast this scenario with that of a second item that also had to be maintained, but which was placed temporarily outside the focus of attention. The data showed that in contrast to the attended, tested-first item, this second tested-second item was maintained without the involvement of any detectable sustained activity, neither in broadband EEG, nor in the alpha band specifically.

Third, previous studies on WM have often analyzed alpha power without distinguishing between evoked and induced components, potentially conflating bottom-up and top-down processes (e.g., Barbosa et al., 2021; Kandemir, Wilhelm, et al., 2024; Sauseng et al., 2005). In our study, evoked (see supplementary materials) and induced alpha activity showed distinct temporal and functional characteristics. Evoked alpha power, which was expected to reflect stimulus-locked responses, primarily associated with the initial encoding of WM items in our task, indeed showed robust decoding of both tested-first and tested-second items shortly after stimulus presentation. However, this decoding was transient, with no sustained decoding observed during the later portions of the delay period. Importantly, induced alpha power, which should reflect internally generated, non-phase-locked oscillatory activity, remained the primary, sustained carrier of prioritized information throughout the long delay condition.

There were two more subtle deviations from prior impulse perturbation studies in the present study. The first is that, in the long delay condition, impulse 1 did not reliably reveal decoding of the tested-second item. This may have been caused by the longer delay, during which representations may have deteriorated enough to become undetectable at this stage. However, the tested-second effect at impulse 1 seems generally weak and brief (cf. Wolff et al., 2017, Figure 6A), even with short delays. We therefore treat its absence after a longer delay as not very diagnostic, as it may reflect further attenuation of an already small effect rather than a qualitative failure of the impulse response to reveal latent representations. Indeed, also in the current data, when the tested-second item became relevant later, at impulse 2, it was robustly decodable, as was previously found as well. The second deviation was that following impulse 2, we observed occasional decoding of the tested-first item, although this item was no longer task-relevant after probe 1. One possibility is that this reflects carry-over activity from the first probe, such as residual retrieval or response-related activity, which may temporarily bias population patterns even when the item is no longer needed. Another possibility is (strategic) incomplete dropping. Participants may not fully remove the tested item when overall load is low, leading to slow decay rather than abrupt removal. Yet, also because this effect at impulse 2 was small, we hesitate to draw strong theoretical conclusions from it.

In view of the overall results, one might ask whether alpha oscillations during WM maintenance serve mainly to protect attended representations against interference, or to prepare for associated action, which is thought to be an important function of WM (van Ede, 2020).The present data cannot fully arbitrate this issue. Against the former account, one might wonder why non-prioritized representations would then not need such protection. Against the latter account, one might consider the raw voltage decoding during the long delay interval. Towards the end of this interval, raw voltage decoding steadily increased, likely in anticipation of the probe, when action was needed. By contrast, such an increase was not apparent in the alpha band, suggesting that this preparatory effect has its roots elsewhere. As such, it would seem that further research is needed to more precisely determine the nature of the prioritization afforded by alpha oscillations in WM.

One further hint towards the functional role of alpha oscillations might be had from the temporal generalization that we observed. Initially, through most of the short delay interval, decoding patterns were highly dynamic and time-specific, consistent with rapidly evolving neural codes (Lundqvist et al., 2016; Wolff et al., 2017). After this initial period, we observed a transition from dynamic to stable representations, in both raw voltage and induced alpha power. The emergence of sustained cross-temporal decoding about 1 s post-stimulus suggests that prolonged maintenance may depend on temporally stable attractor-like states (Murray et al., 2017; Spaak et al., 2017), and may reflect a shift from perceptual encoding to abstract mnemonic representations over time (Kamiński & Rutishauser, 2020; Stokes, 2015). These results support mixed-state models of WM, in which initially dynamic activity gradually turns into a stable coding pattern over the course of the delay, rather than both codes coexisting throughout (Murray et al., 2017).

Overall decoding of tested-second items, or rather the lack thereof, might speculatively be related to this change in representational format over time. Tested-second item decoding fell back to baseline, or at least close to it, within 1.5 s across all measures. Possibly, the representational change that tested-first items go through is mirrored by a transition to a activity-quiescent state for tested-second items; a moment at which unattended information may lose active status. It is also interesting to note that this purported transition occurs when computational models of STSP would suggest that short-term synaptic traces that can sustain WM representations reach the end of their lifespan (Mongillo et al., 2008). Nevertheless, it should be noted that in spite of this convergence the possibility remains that this transition is one towards some other kind of ongoing activity that is undetectable by EEG (e.g., due to the involvement of other brain regions that might be less accessible due to cortical folding; Christophel et al., 2018).

To conclude, our results reveal that sustained neural representations in the alpha band during the delay interval are not uniformly present across all memorized items but instead selectively track the attentional prioritization of WM contents. Specifically, while early decoding accuracy for both prioritized (tested-first) and deprioritized (tested-second) items was evident across raw voltage and evoked alpha power, these signals rapidly diminished over time and failed to support sustained maintenance, particularly for deprioritized (tested-second) items. By contrast, induced alpha power revealed temporally extended and selectively robust decoding of the prioritized item, especially in the long delay condition. Importantly, deprioritized items did not exhibit similarly sustained decoding, suggesting that induced alpha oscillations do not support the general maintenance of all memory contents, but are instead selectively engaged by prioritized representations. These findings suggest that alpha oscillations serve as a neural substrate for attentional prioritization in WM, rather than a generic mechanism for maintaining all stored information therein.

## Declarations

### Data and code availability

All data, analysis scripts and results are available at the Open Science Framework: osf.io/4tejh. The repository is currently private for peer review and will be made publicly available upon acceptance of manuscript.

### Author contributions

Yuanyuan Weng: Data curation, methodology, visualization, software, formal analysis, funding acquisition, investigation, writing – original draft, writing – review and editing.

Jelmer P. Borst: Conceptualization, methodology, formal analysis, supervision, funding acquisition, writing – review and editing.

Elkan G. Akyürek: Conceptualization, methodology, formal analysis, supervision, funding acquisition, writing – review and editing.

## Footnotes

1 The term “activity-silent” state refers to memory retention primarily through short-term synaptic changes rather than sustained neural firing. However, this does not imply a complete absence of neural activity, such as more subtle, intermittent neural firing that would by and of itself be insufficient to maintain memoranda in WM. Hence, the term “activity-quiescent” may be more appropriate.

2 A selective role of alpha oscillations might also comprise priority-dependent refreshing of memoranda. Importantly, this still implies that this mechanism is not applied (equally) to deprioritized items.

