## Supplementary Materials for "Sustained alpha oscillations serve attentional prioritization in working memory, not maintenance"

#### S1. Evoked alpha power decoding

#### S1.1 Delay interval evoked alpha power decoding

As indicated in the main text, we also analyzed evoked alpha power to characterize externally driven neural processes. In the short delay condition, the decoding results for evoked alpha power closely mirrored those observed in the raw voltage time-course analysis. Both the tested-first (Figure S1A, left, 12-752 ms, $p < 0.001$, one-sided, corrected) and tested-second items (Figure S1A, left, 2-536 ms, $p < 0.001$, one-sided, corrected) were decodable, with decoding accuracy peaking shortly after item 2 onset and gradually declining during the maintenance period. Time window of interest analyses showed that in the early time window, both tested-first (Figure S1A, right, $p < 0.001$, one-sided) and tested-second items (Figure S1A, right, $p < 0.001$, one-sided) were decodable. In the late time window, decoding accuracy was much reduced overall, but remained significantly above chance for both items (Figure S1A, right, all $p < 0.001$, one-sided).

In the long delay condition, evoked alpha power reflected both the tested-first (Figure S1B, left, 30-372 ms, $p = 0.001$, and 1640-2616 ms, $p = 0.005$, both one-sided, corrected) and tested-second item (Figure S1B, left, 4-428 ms, $p < 0.001$, one-sided, corrected) immediately after stimulus presentation, similar to the raw voltage decoding results. Time window analyses showed that in the early time window (100-400 ms), both tested-first and tested-second items were decodable (Figure S1B, right, all $p < 0.001$, one-sided), while in the late time window (2400-3000 ms), only the tested-first item remained decodable (Figure S1B, right, tested-first: $p = 0.032$; tested-second: $p > 0.05$, one-sided).

We also compared decoding accuracy across short and long delay conditions. In the late time window of interest, evoked alpha power decoding accuracy was significantly stronger for the short delay than the long delay for both the tested-first item ($t$ (27) = -3.19, $p = 0.004$, *Cohen's d* = -0.60) and the tested-second item ($t$ (27) = -3.01, $p = 0.006$, *Cohen's d* = 0.57). No further comparisons were significant (all $p > 0.10$).

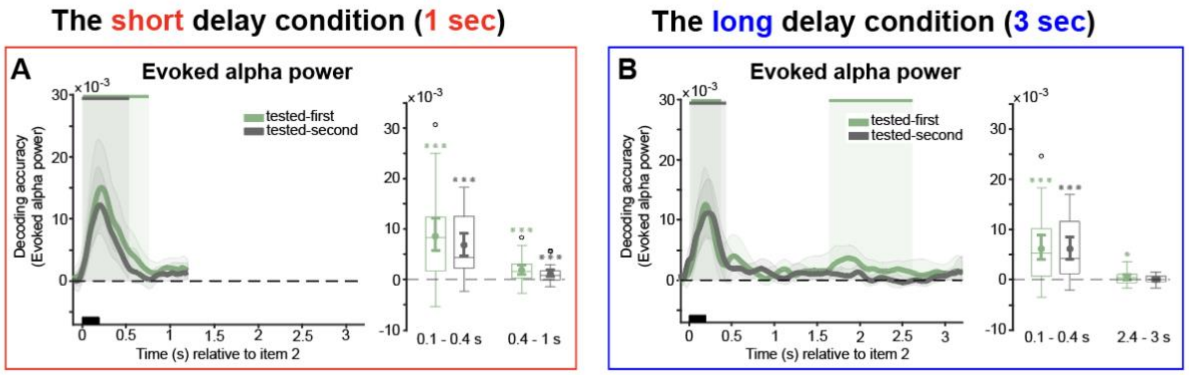


**Figure S1.** Evoked alpha power time course decoding during delay intervals. A, the short delay condition. B, the long delay condition. Upper solid bars and corresponding shading indicate significant decoding areas (one-sided, $p < 0.05$). Error shading represents 95% CI of the mean. Boxplots illustrate decoding accuracy for tested-first and tested-second items across the two selected time windows of interest in the short delay condition (100-400 ms and 400-1000 ms) and the long delay condition (100-400 ms and 2400-3000 ms). The middle line within each box represents the median, while the box itself spans the 25th to 75th percentiles. Whiskers extend to 1.5 times the interquartile range, and individual dots indicate extreme values. Mean decoding accuracy is marked by filled circles, with vertical error bars showing the 95% CI. Asterisks indicate significant decoding (*, $p < 0.05$; **, $p < 0.01$; ***, $p < 0.001$).

#### S1.2 Impulse 1 evoked alpha power decoding

The results from evoked alpha power time course decoding showed that both the tested-first and tested-second item were decodable from impulse 1 in the short delay condition (Figure S2A, left, tested-first: -20-416 ms, $p < 0.001$, one-sided, corrected; tested-second: -80-148 ms, $p = 0.010$, one-sided, corrected). The analysis of the time window of interest revealed significant decoding for the tested-first item (Figure S2A, right, $p < 0.001$, one-sided), and for the tested-second item (Figure S2A, right, $p = 0.006$, one-sided). Moreover, decoding accuracy was significantly higher for the tested-first than for the tested-second item ($t$ (27) = 2.90, $p = 0.007$, Cohen's d = -0.55).

In the long delay condition, consistent with raw voltage decoding, the tested-first item could be decoded from evoked alpha power (Figure S2B, left, 104-466 ms, $p = 0.001$, one-sided, corrected), while the tested-second item could not. Within the time window of interest, significant decoding was observed for the tested-first item (Figure S2B, right, $p < 0.001$, one-sided), but not for the tested-second item (Figure S2B, right, $p = 0.308$, one-sided). A direct contrast confirmed that there was higher decoding for the tested-first than the tested-second item ($t$ (27) = 3.31, $p = 0.003$, *Cohen's d* = 0.62).

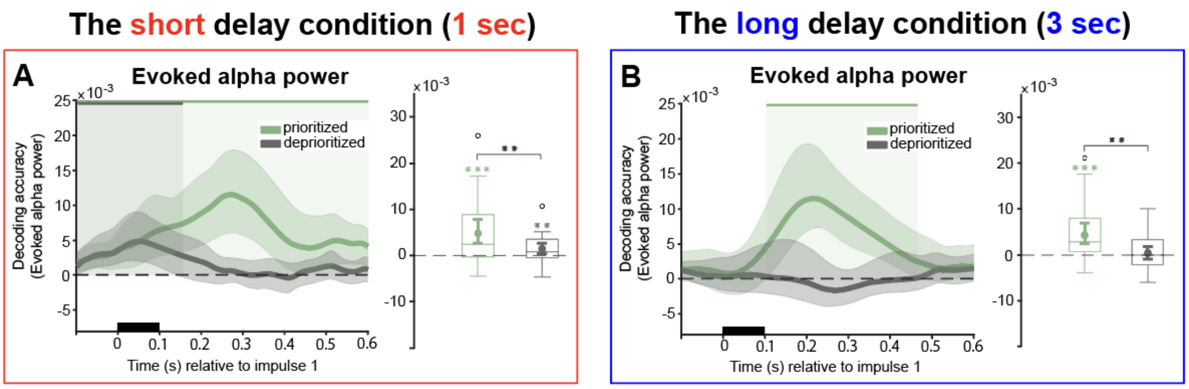


**Figure S2.** Evoked alpha power time course decoding from the first impulse response. A, the short delay condition. B, the long delay condition. Upper solid bars and corresponding shading indicate significant decoding areas (one-sided, $p < 0.05$). Error shading represents 95% CI of the mean. Boxplots illustrate decoding accuracy for tested-first and tested-second items within the time window of interest (100-400 ms). The middle line within each box represents the median, while the box itself spans the 25th to 75th percentiles. Whiskers extend to 1.5 times the interquartile range, and individual dots indicate extreme values. Mean decoding accuracy is marked by filled circles, with vertical error bars showing the 95% CI. Asterisks indicate significant decoding (*, $p < 0.05$; **, $p < 0.01$; ***, $p < 0.001$).

#### S1.3 Impulse 2 evoked alpha power decoding

In the short delay condition, the tested-second item could be decoded from evoked alpha power (Figure S3A, left, 154-600 ms, p < 0.001, one-sided, corrected), with decoding accuracy remaining above chance for a sustained period after impulse presentation. In the time window of interest for evoked alpha power, only the tested-second item could be reliably decoded (Figure 3A, right, p < 0.001, one-sided), while the tested-first item could not (Figure S3A, right, p = 0.250, one-sided). A notable difference emerged in the strength of their representations (t (27) = -3.41, p = 0.002, Cohen's d = -0.64).

A similar pattern emerged in the long delay condition, with reliable decoding of the tested-second item from evoked alpha power (Figure S3B, left, 46-600 ms, $p = 0.001$, one-sided, corrected). In the predefined time widow, significant decoding was again observed only for the tested-second item (Figure S3B, right, $p < 0.001$, one-sided), with a corresponding significant difference between tested-first and tested-second items ($t$ (27) = -3.18, $p = 0.004$, *Cohen's d* = 0.60).

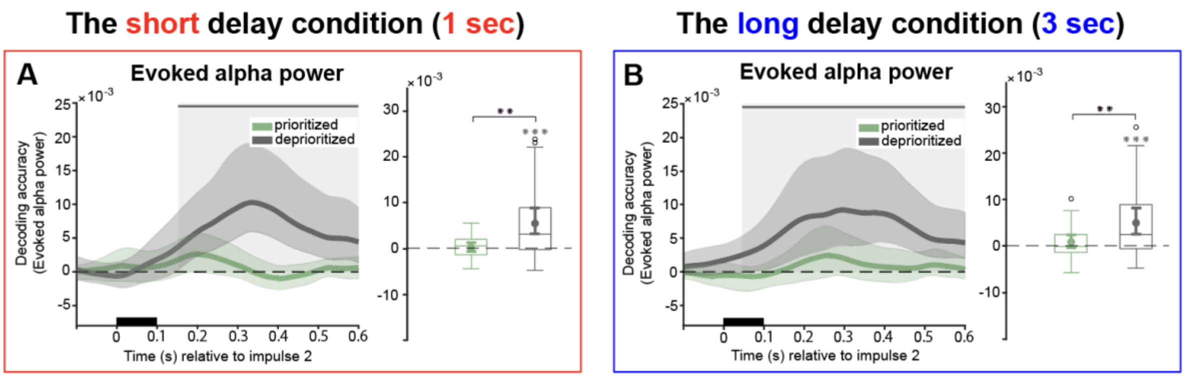


**Figure S3.** Evoked alpha power time course decoding from the second impulse response. A, the short delay condition. B, the long delay condition. Upper solid bars and corresponding shading indicate significant decoding areas (one-sided, $p < 0.05$). Error shading represents 95% CI of the mean. Boxplots illustrate decoding accuracy for tested-first and tested-second items within the time window of interest (100-400 ms). The middle line within each box represents the median, while the box itself spans the $25_{th}$ to $75_{th}$ percentiles. Whiskers extend to 1.5 times the interquartile range, and individual dots indicate extreme values. Mean decoding accuracy is marked by filled circles, with vertical error bars showing the 95% CI. Asterisks indicate significant decoding (*, $p < 0.05$; **, $p < 0.01$; ***, $p < 0.001$).

#### S1.4 Evoked alpha cross-temporal generalization

As might be expected, evoked alpha power reflected mainly stimulus-related activity. Following the onset of memory item 2 in the short delay condition, decoding was strongest along the diagonal, indicating that the representational format changes rapidly over time. Off-diagonal decoding was also present, but it was restricted to a narrow near-diagonal region early in the epoch and did not extend into later portions of the delay. Diagonal decoding was reliably higher than off-diagonal decoding during the same early interval, further supporting that evoked alpha neural activity was mainly dynamic, characterized by rapidly evolving neural representations following the onset of memory item 2, and ending after about half a second (Figure S4A, cluster-based permutation test, cluster-corrected, $p < 0.05$). A similar early dynamic phase was also observed in the long delay condition. Significant decoding was strongest along the diagonal early in the epoch, with off-diagonal generalization remaining spatially limited, rather than extending broadly across time. A complementary diagonal-referenced analysis further showed that off-diagonal decoding was reliably weaker than diagonal decoding, consistent with predominantly dynamic evoked representations rather than a sustained stable code (Figure S4B, cluster-based permutation test, cluster-corrected, $p < 0.05$).

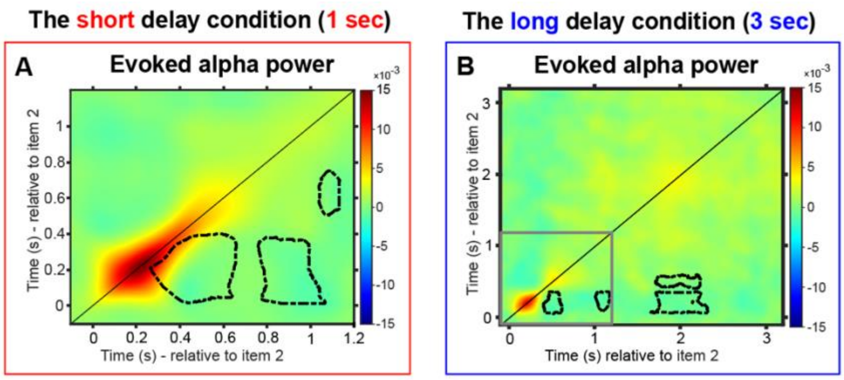


**Figure S4.** Cross-temporal decoding matrices for the prioritized item in evoked alpha power, derived from training and testing on all time-point combinations. A, the short delay condition. B, the long delay condition. Decoding accuracy is represented by color intensity. The black lines indicate time-points of significantly lower decoding relative to both equivalent time-points along the diagonal (same train and test time point, $p < 0.05$, cluster corrected). Note the scale difference illustrated by the grey outline in panel B, which corresponds to the extent of time plotted in panel A.

### S2. Theta-band decoding

#### S2.1 Delay interval evoked theta power decoding

We additionally examined evoked theta power, focusing on whether stimulus-locked activity carried item-specific information. In the short delay condition, both items were decodable (tested-first: Figure S5A, left, 8-602 ms, $p < 0.001$, one-sided, corrected; tested-second: Figure S5A, left, 4-538 ms, $p < 0.001$, one-sided, corrected). Time window of interest analyses further confirmed reliable decoding in the early window (tested-first: Figure S5A, right, $p < 0.001$, one-sided; tested-second: Figure S5A, right, $p < 0.001$, one-sided) and also in the late window (tested-first: Figure S5A, right, $p < 0.01$, one-sided; tested-second: Figure S5A, right, $p < 0.001$, one-sided).

In the long delay condition, time-course decoding was again significant for both items (tested-first: Figure S5C, left, 10-544 ms, $p < 0.001$, one-sided, corrected; tested-second: Figure S5C, left, 12-486 ms, $p < 0.001$, one-sided, corrected). In contrast to the short delay condition, decoding did not extend to the late window: While both items were decodable in the early window (tested-first: Figure S5C, right, $p < 0.001$, one-sided; tested-second: Figure S5C, right, $p < 0.001$, one-sided), neither item was decodable in the late window (tested-first: Figure S5C, right, $p = 0.655$, one-sided; tested-second: Figure S5C, right, $p = 0.466$, one-sided).

Direct comparisons between delay conditions showed that decoding was consistently stronger in the short delay condition than in the long delay condition. The difference was evident in the early window for both the tested-first item ($t$ (27) = -2.68, $p$ = 0.013, *Cohen's d* = -0.51) and the tested-second item ($t$ (27) = -3.37, $p$ = 0.002, *Cohen's d* = -0.64). It also remained significant in the late window (tested-first: $t$ (27) = -2.98, $p$ = 0.006, *Cohen's d* = -0.56; tested-second: $t$ (27) = -3.35, $p$ = 0.002, *Cohen's d* = -0.63).

**S2.2 Delay interval induced theta power decoding**

We next assessed decoding based on induced theta power. In the short delay condition, induced theta decoding was significant for both items (tested-first: Figure S5B, left, 8-476 ms, $p$ < 0.001, one-sided, corrected; tested-second: Figure S5B, left, 24-394 ms, $p$ < 0.001, one-sided, corrected). Time window of interest analyses indicated robust decoding in the early window (tested-first: Figure S5B, right, $p$ < 0.001, one-sided; tested-second: Figure S5B, right, $p$ < 0.001, one-sided). In the late window (400-1000 ms), decoding was substantially reduced: the tested-first item remained just above chance (Figure S5B, right, $p$ = 0.046, one-sided), whereas the tested-second item was not reliably decodable (Figure S5B, right, $p$ = 0.0971, one-sided).

In the long delay condition, time course decoding was again detectable for both items, but remained confined to an early post-stimulus interval (tested-first: Figure S5D, left, 38-402 ms, $p$ < 0.01, one-sided, corrected; tested-second: Figure S5D, left, 40-476 ms, $p$ < 0.001, one-sided, corrected). In the early window (100-400 ms), decoding reached significance for the tested-first item (Figure S5D, right, $p$ = 0.012, one-sided) and for the tested-second item (Figure S5D, right, $p$ < 0.001, one-sided). Importantly, neither item showed reliable decoding in the late window (2400-3000 ms; both $p$ > 0.10).

We also tested for condition differences in the predefined windows. A reliable short and long difference was observed only for the tested-first item in the early window (100-400 ms), with higher decoding accuracy in the short than the long delay condition ($t$ (27) = -2.17, $p$ = 0.039, *Cohen's d* = -0.41). All remaining short-long comparisons were not significant (all $p$ > 0.10).

In sum, while evoked theta showed an early decoding pattern comparable to that observed for evoked alpha, induced theta did not reproduce the sustained delay period decoding that characterized induced alpha. Induced theta decoding was primarily observed early and was markedly reduced later in the delay, supporting the idea that the sustained decoding observed in induced alpha power is frequency-specific (see also beta-band decoding below).

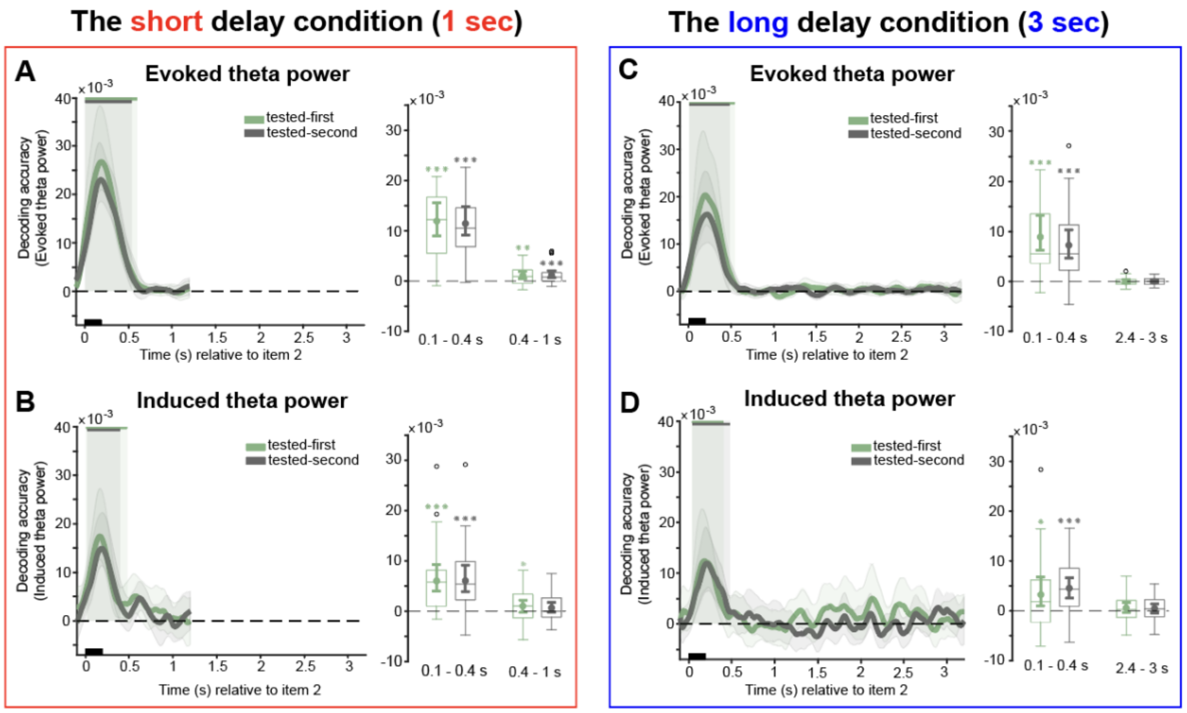


**Figure S5.** Theta-band decoding during delay intervals. A-B, the short delay condition. C-D, the long delay condition. A and C, evoked theta power time course decoding. B and D, induced theta power time course decoding. Upper solid bars and corresponding shading indicate significant decoding areas (one-sided, $p < 0.05$). Error shading represents 95% CI of the mean. Boxplots illustrate decoding accuracy for tested-first and tested-second items across the two selected time windows of interest in the short delay condition (100-400 ms and 400-1000 ms) and the long delay condition (100-400 ms and 2400-3000 ms). The middle line within each box represents the median, while the box itself spans the 25th to 75th percentiles. Whiskers extend to 1.5 times the interquartile range, and individual dots indicate extreme values. Mean decoding accuracy is marked by filled circles, with vertical error bars showing the 95% CI. Asterisks indicate significant decoding (*, $p < 0.05$; **, $p < 0.01$; ***, $p < 0.001$).

### S3. Beta-band decoding

#### S3.1 Delay interval evoked beta power decoding

Evoked beta decoding was significant in an early interval and again later for both tested items (tested-first: Figure S6A, left, 38-170 ms, $p < 0.001$, and 296-384 ms, $p = 0.014$, both one-sided, corrected; tested-second: Figure S6A, left, 52-142 ms, $p < 0.001$, and 266-364, $p = 0.013$, both one-sided, corrected). In the predefined windows, decoding was significant in the early window (100-400 ms; tested-first: Figure S6A, right, $p < 0.001$, one-sided; tested-second:

Figure S6A, right, $p = 0.010$, one-sided). In the late window (400-1000 ms), only the tested-second item reached significance (tested-first: Figure S6A, right, $p = 0.107$, one-sided; tested-second: Figure S6A, right, $p = 0.048$, one-sided).

A similar time-course pattern was observed in the long delay condition, with significant decoding in an early interval and a later interval for both tested items (tested-first: Figure S6C, left, 50-128 ms, $p < 0.001$, and 280-344 ms, $p = 0.019$, both one-sided, corrected; tested-second: Figure S6C, left, 58-190 ms, $p < 0.001$, and 233-404, $p < 0.001$, both one-sided, corrected). In the predefined windows, early-window (100-400 ms) decoding remained significant for both items (tested-first: Figure S6C, right, $p < 0.001$, one-sided; tested-second: Figure S6C, right, $p < 0.001$, one-sided), whereas late-window (2400-3000 ms) decoding was not significant (both $p > 0.10$).

Within the predefined windows, a reliable difference between the short and long delay conditions emerged only for the tested-first item in the late window, indicating higher decoding accuracy in the short than the long delay condition ($t$ (27) = -2.32, $p = 0.028$, *Cohen's d* = -0.44). The remaining comparisons were not significant (all $p > 0.10$).

**S3.2 Delay interval induced beta power decoding**

In the short delay condition, induced beta decoding was detectable but relatively time-limited. For the tested-first item, decoding reached significance in a brief early interval (Figure S6B, left, 76-156 ms, $p = 0.025$, one-sided, corrected). For the tested-second item, decoding was significant both early and near end of the delay (Figure S6B, left, 66-136 ms, $p = 0.003$, and 980-1060, $p = 0.008$, both one-sided, corrected). In the time window of interest analyses, the early window (100-400 ms) did not show evidence for the tested-first item (Figure S6B, right, $p = 0.387$, one-sided), whereas the tested-second item was significant (Figure S6B, right, $p = 0.034$, one-sided). In the late window (400-1000 ms), neither item reached significance (tested-first: Figure S6B, right, $p = 0.053$, one-sided; tested-second: Figure S6B, right, $p = 0.104$, one-sided).

In the long delay condition, the time course analysis did not reveal significant decoding for either item (both $p > 0.10$). Nonetheless, the predefined early window (100-400 ms) showed a significant effect for the tested-first item (Figure S6D, right, $p = 0.012$, one-sided), whereas the tested-second item did not (Figure S6D, right, $p = 1.25$, one-sided). No evidence for decoding was observed in the late window (400-1000 ms) for either item (both $p > 0.10$).

Direct comparisons between short and long delays did not yield significant differences in either the early or late window for either tested item (all $p > 0.10$).

Summarizing, the beta-band analysis showed evoked effects similar to those seen in alpha, but again no evidence for a sustained effect in induced beta power. Taken together with the theta band analyses that showed very similar outcomes, it can be concluded that the sustained effect observed in induced alpha power is specific to that frequency band alone.

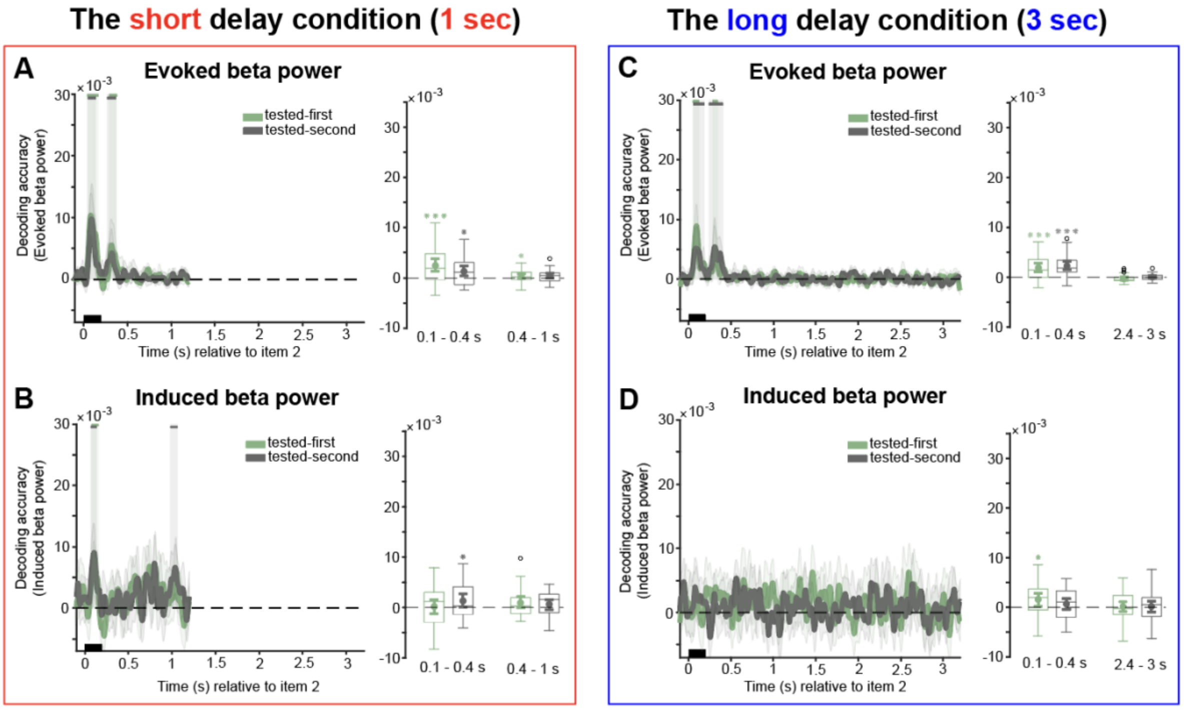


**Figure S6.** Beta-band decoding during delay intervals. A-B, the short delay condition. C-D, the long delay condition. A and C, evoked beta power time course decoding. B and D, induced beta power time course decoding. Upper solid bars and corresponding shading indicate significant decoding areas (one-sided, $p < 0.05$). Error shading represents 95% CI of the mean. Boxplots illustrate decoding accuracy for tested-first and tested-second items across the two selected time windows of interest in the short delay condition (100-400 ms and 400-1000 ms) and the long delay condition (100-400 ms and 2400-3000 ms). The middle line within each box represents the median, while the box itself spans the $25_{th}$ to $75_{th}$ percentiles. Whiskers extend to 1.5 times the interquartile range, and individual dots indicate extreme values. Mean decoding accuracy is marked by filled circles, with vertical error bars showing the 95% CI. Asterisks indicate significant decoding (*, $p < 0.05$; **, $p < 0.01$; ***, $p < 0.001$).

**S4. All-electrodes analysis**

#### S4.1 Delay interval raw voltage decoding

When decoding was performed using all electrodes, above-chance decoding emerged rapidly after the second memory item onset for both tested items in the short delay condition (tested-first: Figure S7A, left, 100-530 ms, $p < 0.001$, one-sided, corrected; tested-second: Figure S7A, left, 110-520 ms, $p < 0.001$, and 540-600 ms, $p = 0.028$, both one-sided, corrected). Consistently, both items were reliably decodable in the predefined early (tested-first: Figure S7A, right, $p < 0.001$, one-sided; tested-second: Figure S7A, right, $p < 0.001$, one-sided) and late (tested-first: Figure S7A, right, $p < 0.001$, one-sided; tested-second: Figure S7A, right, $p < 0.001$, one-sided) windows.

Using the same all electrodes approach in the long delay condition, decoding remained significant shortly after onset for both items in the time-course analysis (tested-first: Figure S7D, left, 90-270 ms, $p = 0.014$, 290-560 ms, $p = 0.009$, 1920-2280 ms, $p = 0.008$, 2300-2580 ms, $p = 0.011$, 2600-3200 ms, $p = 0.003$, all one-sided, corrected; tested-second: Figure S7D, left, 70-480 ms, $p < 0.001$, one-sided, corrected). The time window of interest analyses confirmed significant decoding in the early window for both items (tested-first: Figure S7D, right, $p < 0.001$, one-sided; tested-second: Figure S7D, right, $p < 0.001$, one-sided), and additionally in the late window for the tested-first item (Figure S7D, right, $p = 0.007$, one-sided), whereas the tested-second item (Figure S7D, right, $p = 0.496$, one-sided) was not significant in that late window.

When contrasting the short and long delay conditions with the predefined windows, a significant condition effect was observed only for the tested-second item in the late window, reflecting higher decoding accuracy in the short than the long delay condition ($t$ (27) = -4.60, $p < 0.001$, *Cohen's d* = -0.870), while the other contrasts were not significant (all $p > 0.10$).

#### S4.2 Delay interval evoked alpha power decoding

Using all electrodes, evoked alpha decoding in the short delay condition was robust during the early post-onset period for both items (tested-first: Figure S7B, left, 82-552 ms, $p < 0.001$, one-sided, corrected; tested-second: Figure S7B, left, 34-446 ms, $p < 0.001$, one-sided, corrected). The time window of interest analysis converged with this pattern, showing significant decoding in the early window (100-400 ms) for the tested-first item (Figure S7B, right, $p < 0.001$, one-sided) and the tested-second item (Figure S7B, right, $p = 0.010$, one-sided). In the late window (400-1000 ms), decoding remained reliable for tested-first item (Figure S7B, right, $p = 0.003$, one-sided) but not for tested-second item (Figure S7B, right, $p = 0.165$, one-sided).

In the long delay condition, evoked alpha decoding also reached significance for both items, with effects concentrated in early time ranges (tested-first: Figure S7E, left, 50-128 ms, $p < 0.001$, and 280-344 ms, $p = 0.019$, both one-sided, corrected; tested-second: Figure S7E, left, 58-190 ms, and 238-404 ms $p < 0.001$, both one-sided, corrected). The early window (100-400 ms) decoding was significant for the tested-first item (Figure S7E, right, $p = 0.006$, one-sided) and the tested-second item (Figure S7E, right, $p < 0.010$, one-sided), whereas neither item showed reliable decoding in the late window (400-1000 ms; all $p > 0.10$).

In the window-based short and long delay condition comparisons, a significant condition effect emerged only for tested-first item in the late window, indicating higher decoding accuracy in the short than the long delay condition ($t$ (27) = -2.32, $p = 0.027$, *Cohen's d* = -0.438), while all other window-based contrasts were not significant (all p > 0.10).

**S4.3 Delay interval induced alpha power decoding**

When induced alpha was decoded from all electrodes, the short delay condition revealed a markedly sustained temporal profile for the tested-first item (Figure S7C, left, 62-1196 ms, $p < 0.001$, one-sided, corrected). In contrast, decoding of the tested-second item was comparatively brief and concentrated early in the delay period (Figure S7C, left, 40-278 ms, $p = 0.013$, one-sided, corrected). This difference in temporal extent was also reflected in the time window of interest analysis, which showed reliable early window (100-400 ms) decoding for both items (tested-first: Figure S7C, right, $p = 0.001$, one-sided; tested-second: Figure S7C, right, $p = 0.001$, one-sided), but robust late window (400-1000 ms) decoding only for the tested-first item (Figure S7C, right, $p < 0.001$, one-sided), with the tested-second item reaching significance in the late window evidence (Figure S7C, right, $p = 0.050$, one-sided). Late-window decoding accuracy was higher for the tested-first item than for the tested-second item ($t$ (27) = 2.40, $p = 0.024$, *Cohen's d* = 0.454).

In the long delay condition, the tested-first item again exhibited sustained decoding later in the delay, extending through much of the remaining interval (Figure S7F, left, 804-1204 ms, $p = 0.036$, and 1302-3198 ms, $p < 0.001$, both one-sided, corrected). For the tested-second item, decoding was instead distributed across multiple shorter, separated episodes (Figure S7F, left, 58-464 ms, $p = 0.016$, 814-1260 ms, $p = 0.003$, and 2088-2384 ms, $p = 0.019$, all one-sided, corrected). Early window (100-400 ms) decoding was significant for both items (tested-first: Figure S7F, right, $p = 0.003$, one-sided; tested-second: Figure S7F, right, $p = 0.002$, one-sided), whereas only the tested-first item remained significant in the late window (2400-3000 ms; tested-first: Figure S7F, right, $p < 0.001$, one-sided; tested-second: Figure S7F, right, $p = 0.165$, one-sided). A comparable late-window effect was observed, with decoding accuracy again higher for the tested-first than the tested-second item ($t$ (27) = 2.74, $p = 0.011$, *Cohen's d* = 0.519).

Beyond these effects, we found no additional evidence for differences between the short and long delay conditions in induced alpha power in either the early or late window, and none of the corresponding window-based comparisons in evoked alpha power reached significance (all $p > 0.10$).

In sum, the analyses across all electrodes yielded the same qualitative pattern as the original region-of-interest approach: the key effects reported in the main analyses were preserved, and the overall interpretation remained unchanged. Decoding accuracy was generally lower when using all electrodes than when using 17 posterior electrodes, suggesting that the posterior region of interest (ROI) indeed yields the best decoding of visual WM contents.

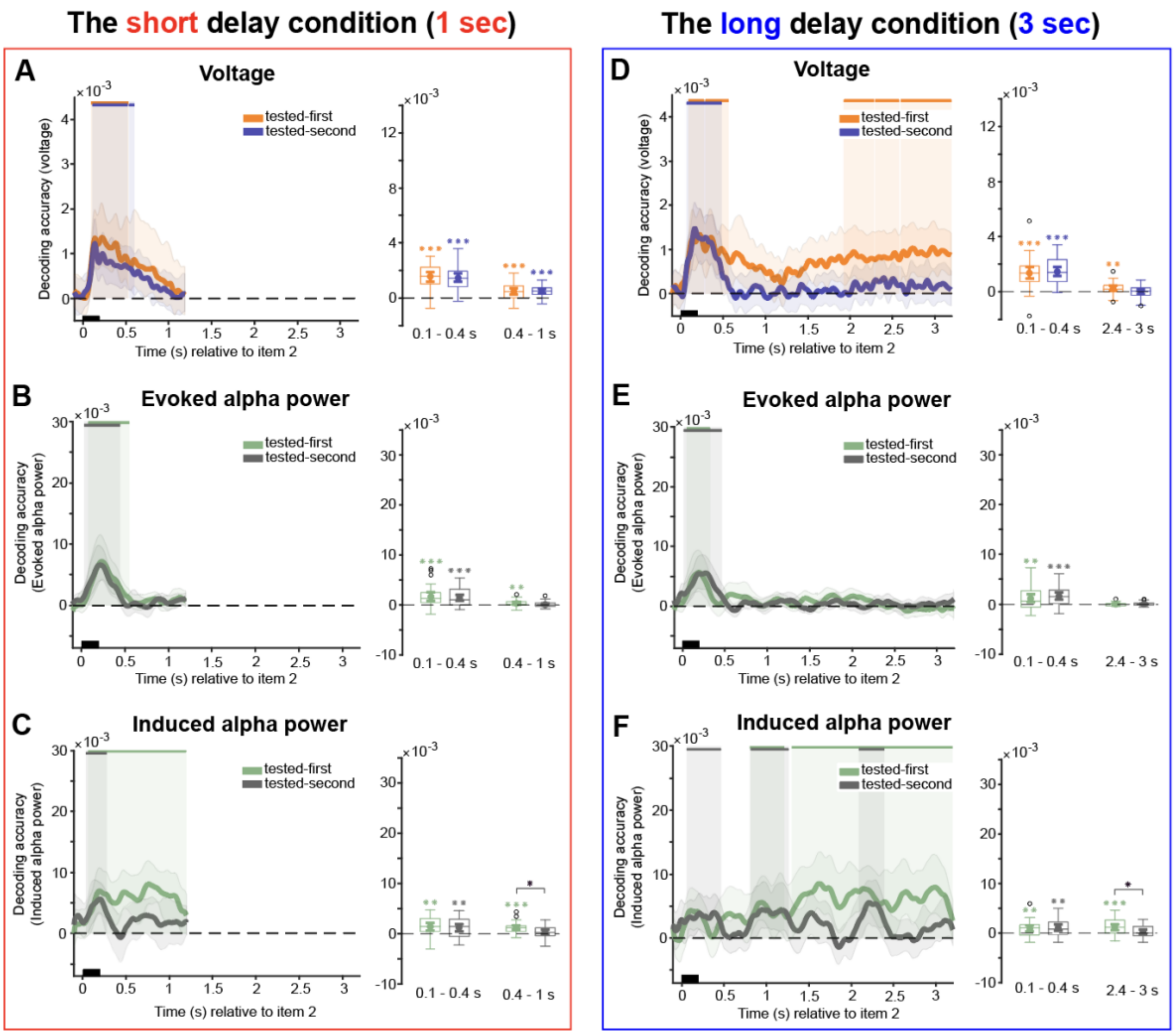


**Figure S7.** Alpha power decoding during delay intervals, using all electrodes. A-C, the short delay condition. D-F, the long delay condition. A and D, raw voltage time course decoding. B and E, evoked alpha power time course decoding. C and F, induced alpha power time course decoding. Upper solid bars and corresponding shading indicate significant decoding areas (one-sided, $p < 0.05$). Error shading represents 95% CI of the mean. Boxplots illustrate decoding accuracy for tested-first and tested-second items across the two selected time windows of interest in the short delay condition (100-400 ms and 400-1000 ms) and the long delay condition (100-400 ms and 2400-3000 ms). The middle line within each box represents the median, while the box itself spans the 25th to 75th percentiles. Whiskers extend to 1.5 times the interquartile range, and individual dots indicate extreme values. Mean decoding accuracy is marked by filled circles, with vertical error bars showing the 95% CI. Asterisks indicate significant decoding (*, $p < 0.05$; **, $p < 0.01$; ***, $p < 0.001$).

### S5. Presentation-order analysis

#### S5.1 Delay interval raw voltage decoding

When trials were re-labeled according to the presentation order of memory items, the short delay condition showed a broadly sustained activity for both items (presented-first: Figure S8A, left, 50-930 ms, $p < 0.001$, one-sided, corrected; presented-second: Figure S8A, left, 70-1040 ms, $p < 0.001$, one-sided, corrected). This pattern was mirrored in the predefined windows: decoding remained significant for both items not only during the early window (100-400 ms; presented-first: Figure S8A, right, $p < 0.001$, one-sided; presented-second: Figure S8A, right, $p < 0.001$, one-sided), but also throughout the late window (400-1000 ms; presented-first: Figure S8A, right, $p < 0.001$, one-sided; presented-second: Figure S8A, right, $p < 0.001$, one-sided).

In the long delay condition, the presented-first item showed significant decoding early and additionally at later points in the delay interval (Figure S8D, left, 40-650 ms, $p < 0.001$, 2800-2920 ms, $p = 0.038$, and 3040-3200 ms, $p = 0.024$, all one-sided, corrected). The presented-second item was significantly decodable only in an early period (Figure S8D, left, 70-560 ms, $p < 0.001$, one-sided, corrected). In the time window analysis, early window (100-400 ms) decoding was significant for both items (presented-first: Figure S8D, right, $p < 0.001$, one-sided; presented-second: Figure S8D, right, $p < 0.001$, one-sided), whereas late window (2400-3000 ms) decoding was not (both $p > 0.10$).

A direct contrast between presented-first and presented-second items showed reliable differences in decoding accuracy within the predefined time windows of interest, with lower decoding accuracy for cued items across several measures. For voltage decoding, the presented-first item was decoded less accurately than the presented-second item in the short delay condition, in both the early and late windows (early: t (27) = -8.600, $p < 0.001$, *Cohen's d* = -1.625; late: t (27) = -6.303, $p < 0.001$, *Cohen's d* = -1.191). A similar effect was observed in the long delay condition in the early window (t (27) = -7.871, $p < 0.001$, *Cohen's d* = -1.488), whereas the late window comparison was not significant ($p > 0.05$)

Direct comparisons between short and long delay conditions mirrored these patterns. In the early window, the condition contrast was not significant for the presented-first item ($p = 0.281$), but was reliable for the presented-second item, with higher decoding accuracy in the short delay than the long delay ($t$ (27) = -2.697, $p = 0.012$, *Cohen's d* = -0.510). In the late window, both items showed robust condition effects, again indicating higher decoding accuracy in the short relative to the long delay condition (presented-first: $t$ (27) = -4.10, $p < 0.001$, *Cohen's d* = -0.775; presented-second: $t$ (27) = -7.37, $p < 0.001$, *Cohen's d* = -1.393).

**S5.2 Delay interval evoked alpha power decoding**

In the short delay condition, both items were reliably decodable in the evoked alpha signal when trials were analyzed by presentation order. Significant decoding was observed for the presented-first item (Figure S8B, left, 12-552 ms, $p < 0.001$, one-sided, corrected) and for the presented-second item (Figure S8B, left, 4-1200 ms, $p < 0.001$, one-sided, corrected). Window-based analyses were consistent with these time-course effects: decoding was significant for both items in the early window (100-400 ms; presented-first: Figure S8B, right, $p < 0.001$, one-sided; presented-second: Figure S8B, right, $p = 0.010$, one-sided), and in the late window (400-1000 ms; presented-first: Figure S8B, right, $p < 0.001$, one-sided; presented-second: Figure S8B, right, $p < 0.001$, one-sided).

In the long delay condition, decoding was again significant early in the time course for both items (presented-first: Figure S8E, left, 18-510 ms, $p < 0.001$, one-sided, corrected; presented-second: Figure S8E, left, 22-726 ms, $p < 0.001$, one-sided, corrected), and both items remained significant in the early window (100-400 ms; presented-first: Figure S8E, right, $p < 0.001$, one-sided; presented-second: Figure S8E, right, $p < 0.001$, one-sided). By contrast, decoding in the late window (2400-3000 ms) was not reliable for the presented-first item (Figure S8E, right, $p = 0.460$, one-sided), whereas only the presented-second item reached significance (Figure S8E, right, $p = 0.045$, one-sided).

For the evoked alpha power, presented-first item again showed reduced decoding, relative to presented-second item, in the short delay condition (early: $t\ (27) = -3.959$, $p < 0.001$, *Cohen's d* = -0.748; late: $t\ (27) = -2.895$, $p = 0.007$, *Cohen's d* = -0.547). In the long delay condition, the early window difference remained significant ($t\ (27) = -4.263$, $p < 0.001$, *Cohen's d* = -0.806), but in the late window the contrast was not significant ($p > 0.05$). Critically, the short and long delay conditions contrast was significant in the late window for the presented-first item, showing higher decoding accuracy in the short than the long delay condition ($t\ (27) = -3.46$, $p = 0.002$, *Cohen's d* = -0.655), whereas all other contrasts did not provide evidence for a reliable condition difference (all $p > 0.10$).

**S5.3 Delay interval induced alpha power decoding**

In the short delay condition, induced alpha decoding was significant for both items when analyzed in presentation order. The presented-first item showed significant decoding in two separated intervals (presented-first: Figure S8C, left, 18-322 ms, $p = 0.003$, and 546-1200 ms, $p < 0.001$, both one-sided, corrected), while the presented-second item was robustly decodable across a sustained time range (Figure S8C, left, 60-1200 ms, $p < 0.001$, one-sided, corrected). Consistent with these time course effects, decoding was significant in both predefined windows. In the early window (100-400 ms), decoding was significant for both items (presented-first: Figure S8C, right, $p = 0.006$, one-sided; presented-second: Figure S8C, right, $p < 0.001$, one-sided). In the late window, decoding was also significant for both items (presented-first: Figure C right, $p = 0.005$, one-sided; presented-second: Figure S8C, right, $p < 0.001$, one-sided).

In the long delay condition, decoding extended markedly further into the delay interval for both items. The presented-first item showed significant decoding for most of the delay (Figure S8F, left, 32-1824 ms, $p < 0.001$, and 1864-2446 ms, $p = 0.018$, both one-sided, corrected). The presented-second item remained significantly decodable almost throughout the delay as well (Figure S8F, left, 48-3234 ms, $p < 0.001$, one-sided, corrected). In the predefined windows, decoding was significant for both items in the early window (100-400 ms; presented-first: Figure S8F, right, $p < 0.001$, one-sided; presented-second: Figure S8F, right, $p < 0.001$, one-sided), and also in the late window (2400-3000 ms; presented-first: Figure S8F, right, $p = 0.026$, one-sided; presented-second: Figure S8F, right, $p = 0.003$, one-sided).

For induced alpha power in the short delay condition, decoding accuracy was lower for the presented-first item than the presented-second item in both windows (early: $t\ (27) = -2.406$, $p = 0.023$, *Cohen's d* = -0.455; late: $t\ (27) = -2.573$, $p = 0.016$, *Cohen's d* = -0.486). Comparing items within each delay condition, short and long comparisons did not yield evidence for condition differences in either window, for either item (both $p > 0.10$).

In sum, when re-labeling items by presentation order, decoding in the short delay condition was generally higher for the presented-second item across raw voltage and evoked alpha, consistent with the proximity to the second-item onset. In the long delay condition, decoding for the two presented items was largely comparable in induced activity. Importantly, this pattern differs from the testing order results, suggesting that presentation order did not confound those results.

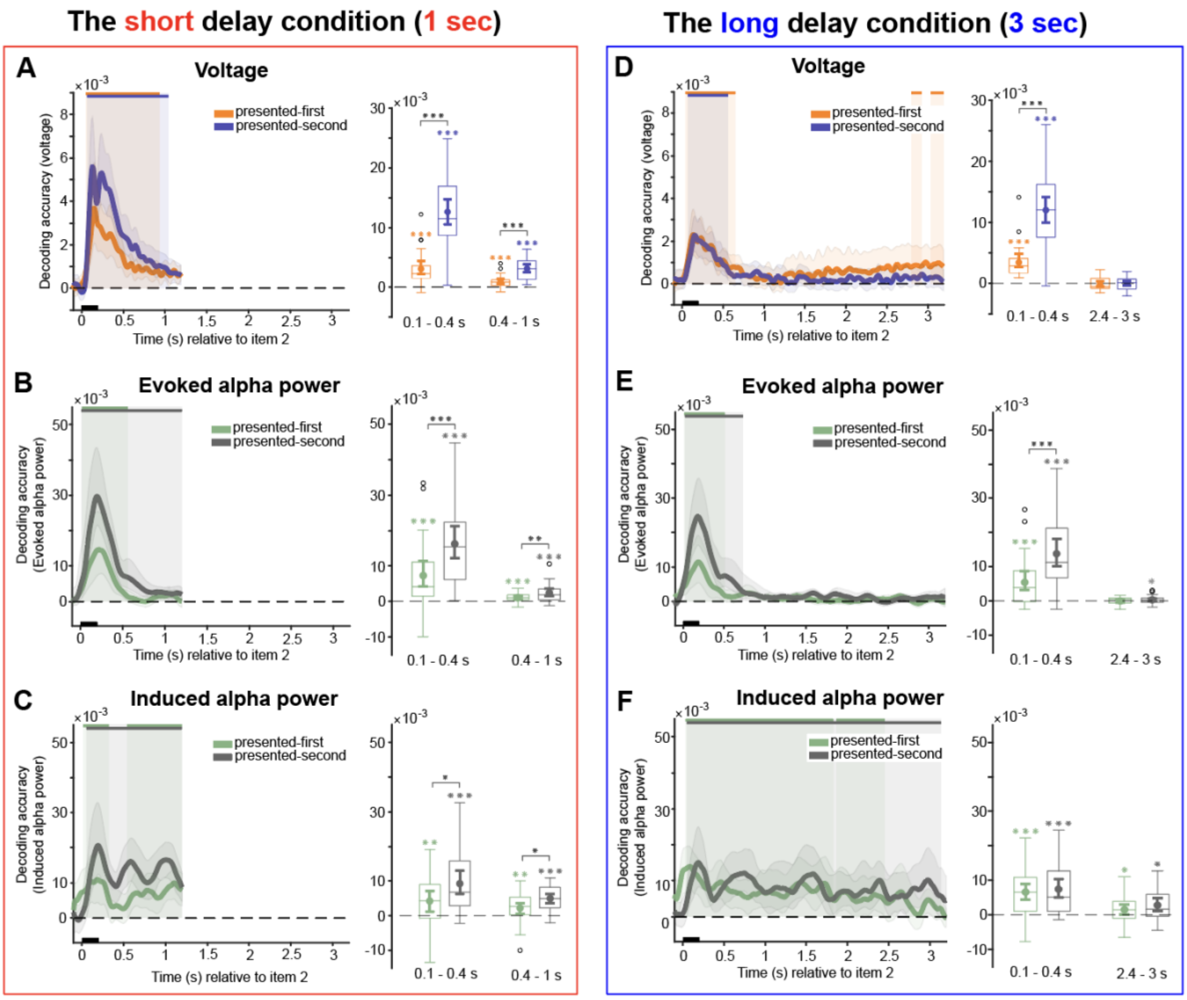


**Figure S8.** Decoding by presentation order during delay intervals. A-C, the short delay condition. D-F, the long delay condition. A and D, raw voltage time course decoding. B and E, evoked alpha power time course decoding. C and F, induced alpha power time course decoding. Upper solid bars and corresponding shading indicate significant decoding areas (one-sided, $p < 0.05$). Error shading represents 95% CI of the mean. Boxplots illustrate decoding accuracy for presented-first and presented-second items across the two selected time windows of interest in the short delay condition (100-400 ms and 400-1000 ms) and the long delay condition (100-400 ms and 2400-3000 ms). The middle line within each box represents the median, while the box itself spans the $25_{th}$ to $75_{th}$ percentiles. Whiskers extend to 1.5 times the interquartile range, and individual dots indicate extreme values. Mean decoding accuracy is marked by filled circles, with vertical error bars showing the 95% CI. Asterisks indicate significant decoding (*, $p < 0.05$; **, $p < 0.01$; ***, $p < 0.001$).
